# Phylogenomic discordance suggests polytomies along the backbone of the large genus *Solanum*

**DOI:** 10.1101/2021.03.25.436973

**Authors:** Edeline Gagnon, Rebecca Hilgenhof, Andrés Orejuela, Angela McDonnell, Gaurav Sablok, Xavier Aubriot, Leandro Giacomin, Yuri Gouvêa, Thamyris Bragionis, João Renato Stehmann, Lynn Bohs, Steven Dodsworth, Christopher Martine, Péter Poczai, Sandra Knapp, Tiina Särkinen

## Abstract

**Premise of the study:** Evolutionary studies require solid phylogenetic frameworks, but increased volumes of phylogenomic data have revealed incongruent topologies among gene trees in many organisms both between and within genomes. Some of these incongruences indicate polytomies that may remain impossible to resolve. Here we investigate the degree of gene-tree discordance in *Solanum,* one of the largest flowering plant genera that includes the cultivated potato, tomato, and eggplant, as well as 24 minor crop plants.

**Methods:** A densely sampled species-level phylogeny of *Solanum* is built using unpublished and publicly available Sanger sequences comprising 60% of all accepted species (742 spp.) and nine regions (ITS, *waxy*, and seven plastid markers). The robustness of this topology is tested by examining a full plastome dataset with 140 species and a nuclear target-capture dataset with 39 species of *Solanum* (Angiosperms353 probe set).

**Key results:** While the taxonomic framework of *Solanum* remained stable, gene tree conflicts and discordance between phylogenetic trees generated from the target-capture and plastome datasets were observed. The latter correspond to regions with short internodal branches, and network analysis and polytomy tests suggest the backbone is composed of three polytomies found at different evolutionary depths. The strongest area of discordance, near the crown node of *Solanum,* could potentially represent a hard polytomy.

**Conclusions:** We argue that incomplete lineage sorting due to rapid diversification is the most likely cause for these polytomies, and that embracing the uncertainty that underlies them is crucial to understand the evolution of large and rapidly radiating lineages.

---

Recent advances in high-throughput sequencing have provided larger molecular datasets, including entire genomes, for reconstructing evolutionary relationships (e.g. Ronco et al., 2021). Considerable progress has been made since the publication of the first molecular-based classification of orders and families of flowering plants (APG, 1998), with one of the most recent examples including a phylogenetic tree of the entire Viridiplantae based on transcriptome data from more than a thousand species (One Thousand Plant Transcriptomes Initiative, 2019). Whilst large datasets have strengthened our understanding of evolutionary relationships and classifications across the Tree of Life, several of them have demonstrated repeated cases of persistent topological discordance across key nodes in birds (Suh et al., 2015; Suh, 2016), mammals (Morgan et al., 2013; Romiguier et al., 2013; Simion et al., 2017), amphibians (Hime et al., 2021), plants (Wickett et al., 2014; One Thousand Plant Transcriptomes Initiative, 2019), and fungi (Kuramae et al., 2006). Whereas previous expectations were that these “soft polytomies” would be improved with the addition of more data, their persistence after addition of more taxonomic and molecular data have led some authors to suggest that they actually represent “hard polytomies”, i.e., extremely rapid divergence events of three or more lineages at the same time or reticulate evolution due to species hybridisation and/or introgression. In an era where obtaining genome-wide sampling of species for phylogenetic reconstruction has become mainstream, the question about whether persistent topological discordance can be resolved with more data or whether they reflect complex biological realities (Jeffroy et al., 2006; Philippe et al., 2011) is becoming increasingly common.

Discordance in phylogenetic signal can be due to three general classes of effects (Wendel and Doyle, 1998): (1) technical causes such as gene choice, sequencing error, model selection, or poor taxonomic sampling (Philippe et al., 2011, 2017); (2) organism-level processes such as rapid or convergent evolution, rapid diversification, incomplete lineage sorting (ILS), or horizontal gene transfer (Degnan and Rosenberg, 2009), and (3) gene and genome-level processes such as interlocus interactions and concerted evolution, intragenic recombination, use of paralogous genes for analysis, and/or non-independence of sites used for analysis. Together, these biological and non-biological processes can lead to conflicting phylogenetic signals between different loci in the genome and hinder the recovery of the evolutionary history of a group (Degnan and Rosenberg, 2009). Consequently, careful assessment of phylogenetic discordance across mitochondrial, plastid, and nuclear datasets is critical for understanding realistic evolutionary patterns in a group, as traditional statistical branch support measures fail to reflect topological variation of the gene trees underlying a species tree (Liu et al., 2009; Kumar et al., 2012).

Here we explore the presence of topological discordance in nuclear and plastome datasets of the large and economically important angiosperm genus *Solanum* L. (Solanaceae), which includes 1,228 accepted species and several major crops and their wild relatives, including potato, tomato and brinjal eggplant (aubergine), as well as at least 24 minor crop species (Solanaceaesource.org, Nov. 2020). Building a robust species-level phylogeny for *Solanum* has been challenging because of the sheer size of the genus, and because of persistent poorly resolved nodes along the phylogenetic backbone. Bohs (2005) published the first plastid phylogenetic analysis for *Solanum* and established a set of 12 highly supported clades based on her strategic sampling of 112 species (9% of the total species number in the genus), spanning morphological and geographic variation. As new studies have emerged with increased taxonomic and genetic sampling (e.g., Levin et al., 2006; Weese and Bohs, 2007; Stern et al., 2011; Särkinen et al., 2013; Tepe et al., 2016), the understanding of overall phylogenetic relationships within *Solanum* has evolved to recognise three main clades: (1) the Thelopodium clade containing three species sister to the rest of the genus, (2) Clade I containing c. 350 mostly herbaceous and non-spiny species (including the Tomato, Petota, and Basarthrum clades that contain the cultivated tomato, potato, and pepino, respectively), and (3) Clade II consisting of c. 900 predominantly spiny and shrubby species, including the cultivated brinjal eggplant (Table 1). The two latter clades are further resolved into 10 major and 43 minor clades (Table 1).

**Table 1.**
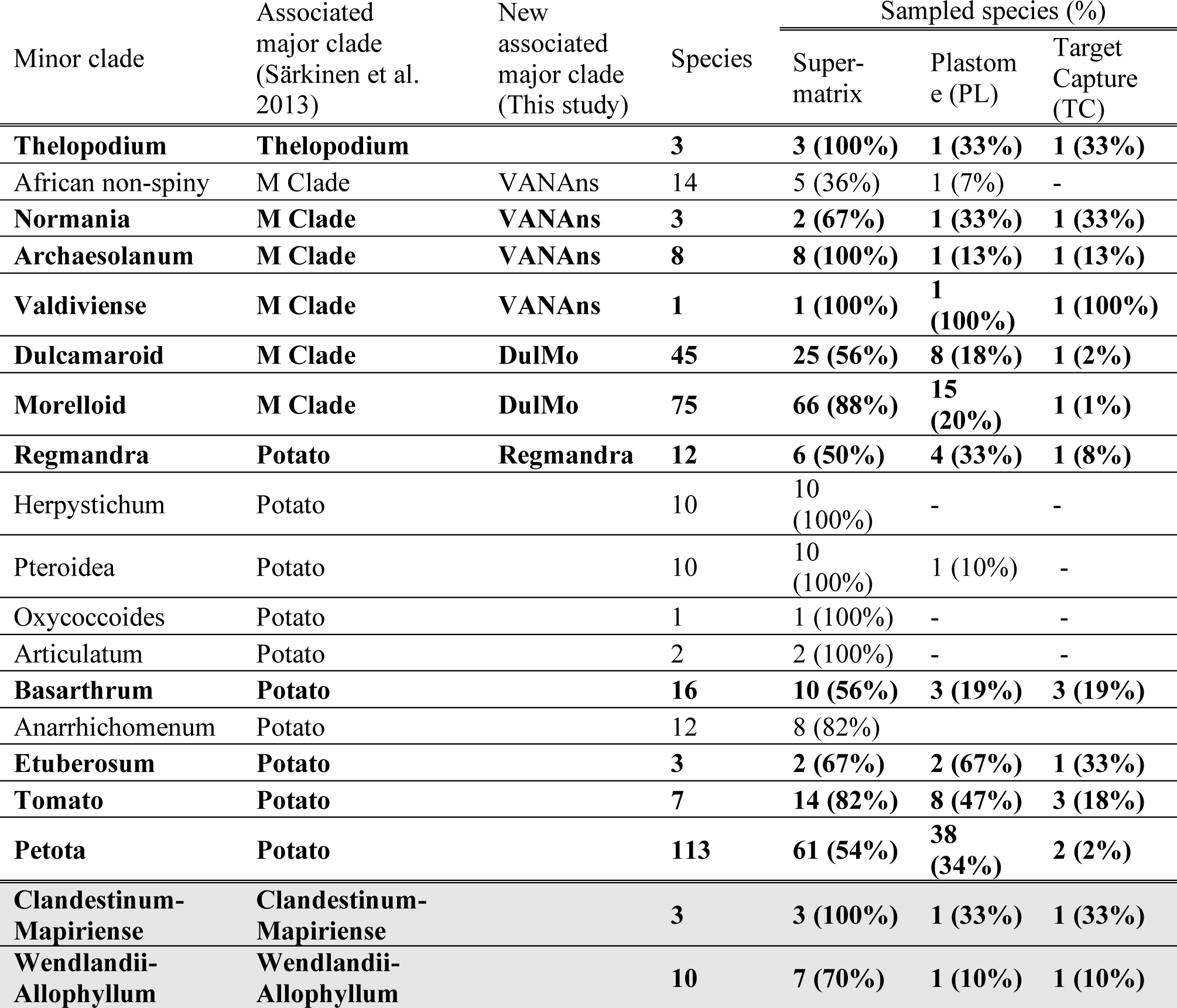
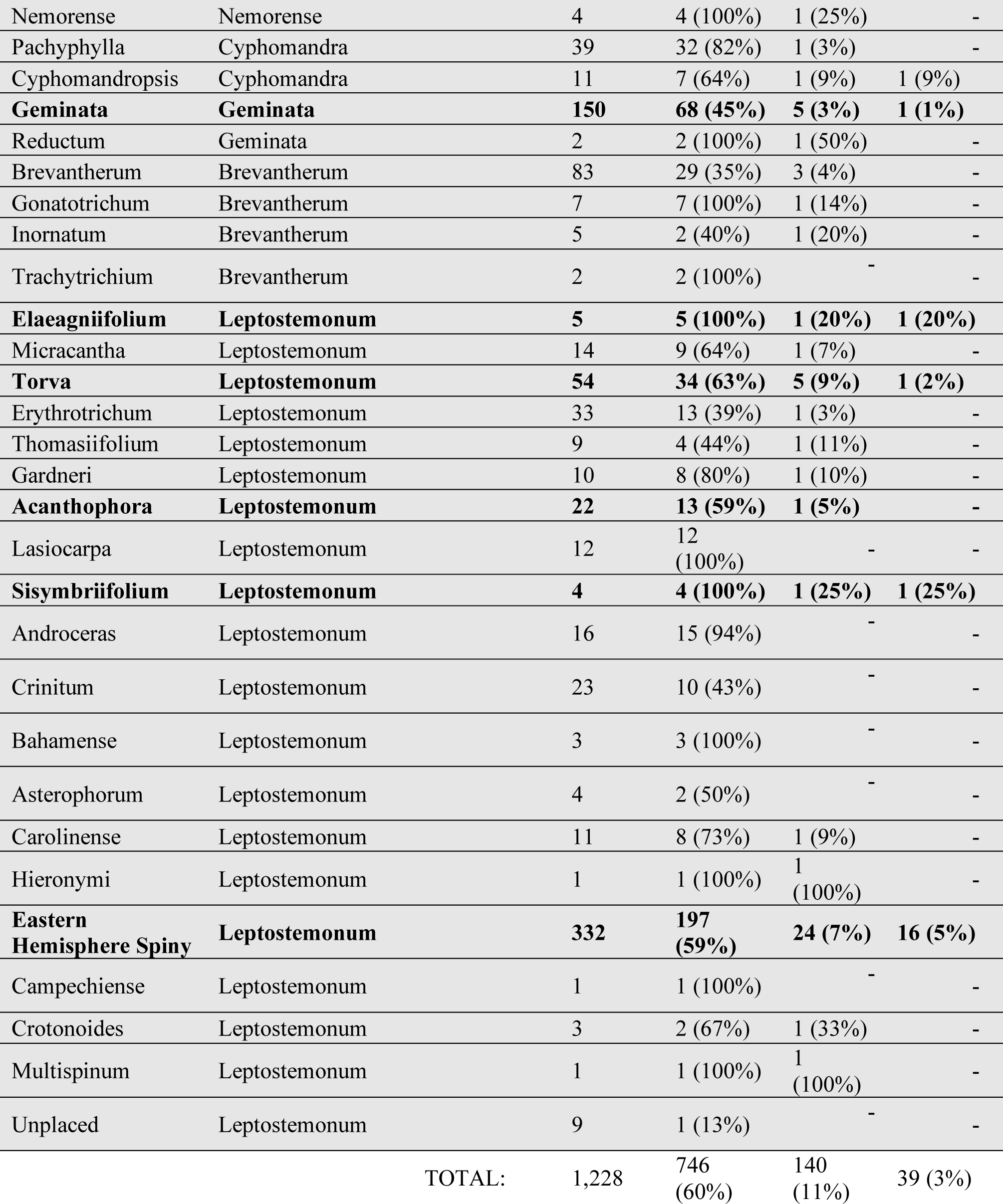
Number of species and taxon sampling across major and minor clades of Solanum. Clades are based on groups identified in previous molecular phylogenetic studies (Bohs, 2005; Weese and Bohs, 2007; Stern et al., 2011; Stern and Bohs, 2012; Särkinen et al., 2013; Tepe et al., 2016). Species number for each clade is based on current updated taxonomy in the SolanaceaeSource database. The 19 clades sampled in the pruned trees for the principal coordinate analysis in this study are in bold. New associated major clade names are given where applicable. Rows shaded in gray represent major and minor clades belonging to Clade II. The Eastern Hemisphere Spiny clade (EHS, formerly known as Old World spiny clade) comprises almost all the spiny solanums occurring in the eastern hemisphere.

Despite these advancements, phylogenetic relationships between many of the major clades of *Solanum* have remained poorly resolved, mainly due to limitations in taxon and molecular marker sampling. The most recent genus-wide phylogenetic study by Särkinen et al. (2013), based on seven markers (two nuclear and 5 plastid) and fewer than half (34%) of the species of *Solanum,* failed to resolve the relationships between major clades, especially within Clade II and the large component Leptostemonum clade, which includes the Old World spiny clade, comprising almost all spiny *Solanum* species that occur in the eastern hemisphere. To reduce colonial connotations associated with this name, we hereafter refer to this clade as the Eastern Hemisphere Spiny clade (EHS; Table 1).

To gain a better understanding of the evolutionary relationships of *Solanum,* we built a new Sanger supermatrix that included 60% of the species of the genus and compared the phylogenetic relationships obtained with the Sanger supermatrix with genus-wide plastid (PL) and nuclear target-capture (TC) phylogenomic datasets. We ask: (1) Does a significant increase in taxon sampling of the supermatrix dataset lead to significant changes in the circumscription of major and minor clades in *Solanum*?; (2) Does increased gene sampling in both plastome and nuclear data resolve previously identified polytomies between major clades?; (3) Is there evidence of discordance within and between genomic datasets?; and (4) Are areas of high discordance in the *Solanum* phylogeny better represented by polytomies rather than bifurcating nodes? Comparison of the topologies from the different datasets, and results from discordance analyses, a filtered supertree network, and polytomy tests lead us to suggest that some of the soft polytomies of *Solanum* might be hard polytomies caused by rapid speciation and diversification coupled with ILS. We discuss the consequences that such an interpretation has for investigating the biogeography and morphological trait evolution across the economically important genus.

## MATERIAL AND METHODS

### Taxon sampling

A Sanger sequence supermatrix was generated including all available sequences from GenBank related to the genus *Solanum* for nine regions: the nuclear ribosomal internal transcribed spacer (ITS), low-copy nuclear region *waxy* (i.e., GBSSI), two protein-coding plastid genes *matK* and *ndhF*, and five non-coding plastid regions (*ndhF-rpl32*, *psbA-trnH*, *rpl32-trnL*, *trnS-G*, and *trnT-L*). Only vouchered and verified samples were utilized. All sequences were blasted against target regions in USEARCH v.11 (Edgar, 2010). Taxon names were checked against SolanaceaeSource synonymy (solanaceaesource.org, Nov. 2020) and duplicate sequences belonging to the same species were pruned out to retain a single individual per taxa. A total of 817 Sanger sequences were generated and added to the matrix, adding 129 previously unsampled species and new data for 257 species (Appendix S1; see the Supplementary Data with this article). Final species sampling across major and minor clades of *Solanum* varied from 13-100%, with 742 species of *Solanum* (60% of the 1,228 currently accepted species, Nov 2020; Table 1). Four species of *Jaltomata* Schltdl. were used as an outgroup (Appendix S1).

To assess phylogenetic discordance within *Solanum*, a set of species was selected for the phylogenomic study to represent all 10 major and as many of the 43 minor clades of *Solanum* as possible (Table 1), as well as the outgroup *Jaltomata*. The final sampling included 151 samples for the plastome (**PL**) dataset (140 *Solanum* species; Table 1 and Appendix S2) and 40 samples for the target-capture (**TC**) dataset (39 *Solanum* species; Table 1 and Appendix S3). For the PL dataset, 86 samples were sequenced using low-coverage genome skimming, and the remaining samples were downloaded from GenBank (Nov 2019). For the TC dataset, 12 samples were sequenced as part of the Plant and Fungal Trees of Life project (Baker et al., 2021) using the Angiosperms353 bait set (Johnson et al., 2019). In addition, 17 sequences were added from an unpublished dataset provided by A. McDonnell and C. Martine. Sequences for the remaining 12 samples were extracted from the GenBank SRA archive using the SRA Toolkit 2.10.7 (https://github.com/ncbi/sra-tools; Appendix S3).

### DNA extraction, library preparation & sequencing

*Supermatrix Sanger sequencings***–** DNA extractions for Sanger sequencing were done using DNeasy plant mini extraction kits (Qiagen, Valencia, California, USA) or the FastDNA kit (MP Biomedicals, Irvine, California, USA). Amplification of *waxy* followed Levin et al. (2005) using two (waxyF with 1171R and 1058F with 2R) or four primer pairs (waxyF with Ex4R, Ex4F with 1171R, 1058F with 3’N, and 3F with 2R). *trnT-L* was amplified with primers a-d and c-f (Taberlet et al., 1991; Bohs and Olmstead, 2001; Bohs, 2004). *ndhF* amplification followed Bohs and Olmstead (1997), *psbA-trnH* followed Sang et al. (1997), *matK* followed Rosario et al. (2019), ITS and *trnS-G* followed Levin et al. (2006), and *rpl32-trnL* and *ndhF-rpl32* followed Miller et al. (2009). Sequencing was carried out on ABI automated sequencers at the University of Utah DNA sequencing facility (Salt Lake City, UT, USA), at the Natural History Museum (London, United Kingdom), and at Myleus Biotecnologia (Belo Horizonte, Brazil). Contigs were visually checked in Sequencher v.4.8 (GeneCodes, Ann Arbor, Michigan, USA) and Geneious Prime 2020.1.1 (https://www.geneious.com). The combined matrix was 10,908 bp long (Appendix S4). The two most densely sampled regions (*trnT-L* and ITS) included 84% and 82% of the sampled species, respectively; *waxy* (54%) and ITS (67%) loci had the most parsimony informative characters (Appendix S4).

*PL and TC datasets–* DNA for high-throughput sequencing was extracted using the low-salt CTAB method (Arseneau et al., 2017) and quantified on a Qubit fluorometer (Thermo Fisher Scientific, USA). Genome skimming was done at the Institute of Biotechnology, University of Helsinki (Finland). A paired-end genomic library was constructed using the Nextera DNA library preparation kit (Illumina, San Diego, CA, USA). Fragment analysis was conducted with an Agilent Technologies 2100 Bioanalyzer using a DNA 1000 chip. Sequencing was performed on an Illumina MiSeq platform from both ends with a read length of 150 bp. DNA extraction, quantification, and sequencing for TC followed Johnson et al. (2019). All PL and TC reads have been submitted to Genbank and the European Nucleotide Archive (Appendices S2-S3).

### Phylogenetic analyses

*Overview of methodological strategy**–*** Ten phylogenetic analyses with different methodological strategies were compared across the supermatrix, PL and TC datasets, to test if the phylogenetic results were robust despite these different choices (e.g., Philippe et al., 2011, 2017; Saarela et al., 2018; Duvall et al., 2020). The Sanger supermatrix analyses based on Maximum Likelihood (ML) and Bayesian inference (BI) were used as a reference to compare results from the PL and TC species trees because the Sanger supermatrix had the most complete taxonomic sampling (Table 2). For the PL dataset, a total of four analysis were compared to test the effect of missing data and sampling on the resulting phylogenies, as well as the effect of different partitioning schemes in IQ-TREE 2 (Table 2; Minh, Schmidt, et al., 2020). For the TC dataset, a total of four analyses were compared to test the effect of the phylogenetic method (ML vs. coalescent methods), missing data, and taxonomic sampling on the resulting phylogenies (Table 2). Full methods for all analyses are described below. All bioinformatic analyses were run either on the Toby-G1 server at the Royal Botanical Gardens of Edinburgh, or the Crop Diversity Server from the James Hutton Institute, in Dundee, Scotland, except for the supermatrix ML analysis.

**Table 2.** Overview of the 10 different analyses conducted across the Sanger supermatrix, plastome (PL) and target capture (TC) datasets. Acronyms indicate how each analysis is referred to in the figures and text. ML = Maximum Likelihood; BI = Bayesian Inference, A353 = Angiosperms353 bait set. See Methods section for full details.

| Dataset | Taxon & genomic sampling | Phylogenetic method | Partitioning scheme | Acronym |
| --- | --- | --- | --- | --- |
| Supermatrix | 746 taxa, 9 loci | ML: RaxML | -- | Supermatrix ML |
|  |  | BI : Beast2 | -- | Supermatrix BI |
| Plastome (PL) | 151 taxa, full + partial plastomes | ML: IQ-TREE2 | Unpartitioned | PL-151-UP |
|  | 151 taxa, full + partial plastomes | ML: IQ-TREE2 | Best-Partition scheme | PL-151-BP |
|  | 125 taxa, full plastomes only | ML: IQ-TREE2 | Unpartitioned | PL-125-UP |
|  | 125 taxa, full plastomes | ML: IQ-TREE2 | Best-Partition scheme | PL-125-BP |
| Target Capture (TC) (A353) | 40 taxa, 338 exons | ML: IQ-TREE2 | -- | TC-min04-ML |
|  | 40 taxa, 338 exons | Coalescent: ASTRAL-III | -- | TC-min04-ASTRAL-III |
|  | 40 taxa, 303 exons | ML: IQ-TREE2 | -- | TC-min20-ML |
|  | 40 taxa, 303 exons | Coalescent: ASTRAL-III | -- | TC-min20-ASTRAL-III |

*Supermatrix dataset*– Sequences were aligned in MAFFT v.7 (Katoh et al., 2005), manually checked, and optimised. Short multi-repeats and ambiguously aligned regions were excluded manually or with trimAl (-gappyout method; Capella-Gutiérrez et al., 2009). Both ML and BI analyses were run on individual loci, as well as on a combined plastid alignment (seven loci in total) to check for topological incongruences, rogue taxa, and misidentified sequences.

Visual checks revealed a small number of clear mis-determinations and/or lab errors. A further 26 samples were removed based on high RogueNaRok scores (Aberer et al., 2013). Nuclear sequence data (ITS and *waxy*) were identified for all known polyploid species (63 species, Appendix S5), and subsequently examined to determine if there were any strong incongruences with the results from the plastid loci. As none were found (Appendices S6-S7), sequences from these species were kept in the final supermatrix analysis.

Maximum likelihood (ML) and Bayesian inference (BI) analyses were run on all nine loci individually and on the combined plastid dataset (seven loci). ML analyses were run in RaxML-HPC v.8.2.12 (Stamatakis, 2014) on XSEDE on CIPRES Science Gateway v.3.3 (Miller et al., 2010), with 10 independent runs based on unique starting trees. The General Time Reversible (GTR) model with CAT (Tavaré and Others, 1986; Stamatakis, 2006) was used for all partitions. A total of 1,000 non-parametric bootstraps were run; bootstrap support (BS) ≥95% was considered strong, 75-94% moderate, and 60-74% weak.

BI analyses were run using Beast v.2.6.3 (Bouckaert et al., 2019), with two parallel runs sampling trees every 10,000 generations. ModelTest-NG (Darriba et al., 2020) was used to find the most suitable nucleotide substitution model for the individual loci and combined plastid loci; JC + G4 was specified for the *ITS* and *trnS-G* regions, GTR+G4 for the *psbA-trnH, trnL-T, rpL32* and *matK* regions, and the GTR+I+G4 model for all other regions, as well as the combined plastid dataset and the full supermatrix dataset. For all analyses, an uncorrelated log normal relaxed clock, birth-death tree prior, and a normally distributed UCLD.mean prior was specified (mean 1, SD=0.3). All runs were checked with Tracer v.1.7.1 (Rambaut et al., 2018) to ensure that adequate effective sample sizes were reached (ESS >200). LogCombiner and TreeAnnotator were used to generate the final maximum credibility tree with a 15% burn-in. Posterior probability (PP) values ≥0.95 were considered strong, and from 0.94 to 0.75 as moderate to weak.

The concatenated ML Sanger supermatrix analysis was run on a concatenated matrix, with the same settings as described above in RaxML. The concatenated BI Sanger supermatrix was analysed partitioning the dataset between ITS, *waxy* and the plastid genes. Modifications to the analysis included a monophyletic constraint on *Solanum*, and four parallel runs that were run for 60 million generations with two chains, sampling trees every 10,000 generations. The ML best tree was used as a starting topology to speed up convergence of the chains.

*PL dataset–* Paired reads from genome skimming were cleaned using BBDuk from the BBTools suite (<u>sourceforge.net/projects/bbmap/</u>; ktrimright=t, k=27, hdist=1, edist=0, qtrim=rl, trimq=20, minlength=36, trimbyoverlap=t, minoverlap=24, and qin=33). Sequence quality was checked with FastQC (Andrews and Others, 2010) and MultiFastQC (Ewels et al., 2016). Plastome assembly was done using de-novo assembly with Fast-Plast v.1.2.6 (https://github.com/mrmckain/Fast-Plast), and reference-guided assembly using GetOrganelle v.1.6.2.e (Jin et al., 2020) with the high-coverage plastome sequence of *S. dulcamara* L. (GenBank KY863443 (Amiryousefi et al., 2018). For GetOrganelle, the following settings were used: -w 0.6, -R 20 -k 85, 95, 105, 127; for Fast-Plast, the Solanales Bow-tie index was used for the assembly. Results from both methods were aligned in Geneious and visually checked to determine consistency. Assembly quality was assessed using the reads identified from the Bow-tie step in the Fast-Plast analysis, which were mapped against the final recovered plastome sequence using BWA (Li and Durbin, 2010). Mean and standard deviation of coverage depth for each base pair was determined by examining the same files in Geneious. Assemblies were annotated using both Chlorobox GeSeq (Tillich et al., 2017) and the “Annotate from database” tool in Geneious using the reference plastid genome of *S. dulcamara*. Results were compared to ensure that start and stop codons for exon boundaries were congruent. Annotated plastomes were submitted to Genbank (Appendix S2). A total of 55 full plastomes were assembled with a mean length of 155,498 bp (max. 156,138 bp, min. 154,715 bp; Appendix S2), and a mean coverage of 158 (min. 22, max. 571; Appendix S2), and 28 partial plastomes (45,398-154,598 bp) with a mean coverage of 29 (min 4, max 96; Appendix S2). All plastomes had a highly conserved quadripartite structure, with no loss, duplication, or expansion of gene families.

Plastomes from this study and those retrieved from Genbank were aligned in Geneious using MAFFT (Katoh et al., 2005), visually checked, and corrected. A copy of the inverted repeat (IRa) was removed prior to phylogenomic analyses, although 1,189 bp were kept at the beginning of the region to be able to extract the gene that spans the boundary between the small single copy (SSC) and IRa region. We then separated the plastome alignment into: (1) 79 protein-coding regions, (2) 15 introns, (3) and 73 intergenic regions. For each dataset, the ambiguously aligned regions and polyA repeats were removed, using visual checks for the exons and intron regions, and the strict mode of trimAl (Capella-Gutiérrez et al., 2009) for the intergenic regions (Appendix S8). Sequences shorter than 25% of the length of the aligned matrix for each region and columns containing >75% of gaps were removed in trimAl (Capella-Gutiérrez et al., 2009) to avoid issues with long branch attraction following Gardner et al. (2021). Two pseudogenes (*ycf*1 and *rps*19) at the junction of IRa and Long Single Copy (LSC) (Amiryousefi et al., 2018), and four intergenic regions with no parsimony informative characters were excluded from the final analysis. All remaining loci alignments were concatenated together for the final PL phylogenetic analyses.

To test for the effect of missing data, two datasets were compared: a matrix with 151 taxa containing all 140 species selected for this study with higher proportion of missing data (147,278 bp long with the second IR removed), and a matrix with 125 samples containing only complete plastid sequences (Appendices S2 and S8).

ML searches were run on all PL datasets in IQ-TREE2 (Minh, Schmidt, et al., 2020) with 1,000 non-parametric bootstraps. Optimal substitution models were determined using –TEST in IQ-TREE2 (Appendix S9). For both PL datasets, topologies from two different partitioning schemes were also compared (unpartitioned vs. best-fit partition scheme based on PartitionFinder; Lanfear et al., 2012) in IQ-TREE 2, to test if accounting for variation in substitution rate amongst loci affected the phylogenetic results. BS values ≥95% were considered strong, 75-94% moderate, and 60-74% weak.

*TC dataset–* Trimmomatic (Bolger et al., 2014) was used to trim reads (TruSeq3-PE- simpleclip.fa:1:30:6, LEADING:30, TRAILING:30, SLIDINGWINDOW:4:30, MINLEN:36). Read quality was checked with FastQC (Andrews and Others, 2010) and MultiFastQC (Ewels et al., 2016). Over-represented repeat sequences were removed with CutAdapt (Martin, 2011). HybPiper (Johnson et al., 2016) was used to produce reference-guided *de novo* assembles using the reference provided by Johnson et al. (Johnson et al., 2019). Putative paralogs were identified using the HybPiper script “paralog_retriever.py”. Phylogenies were generated for all 45 loci for which paralog warnings were found using MAFFT (Katoh et al., 2005) and FastTree (Price et al., 2010). Five loci were deleted and several taxa whose paralogs caused paraphyly of clades were excluded from 27 loci (one to seven taxa per loci). A single gene (g5299) presented a clear duplication event and was divided into two separate matrices for downstream analyses.

Default HybPiper settings were used for all but three samples (*S. betaceum* Cav., *S. valdiviense* Dunal, and *S. etuberosum* Lindl.), for which the coverage cutoff was reduced from eight to four to maximise recovery of target genes. One sample (*S. terminale* Forssk.) was excluded due to poor sequence quality. Only the exon dataset was analysed in downstream phylogenomic analyses, because the transcriptome dataset showed large differences in the recovered flanking regions of target loci between samples, likely due to post-transcriptional splicing and editing of messenger RNA. The HybPiper script “fasta_merge.py” was used to concatenate all genes together and produce a partition file. In summary, an average of 289 genes per sample were recovered for the TC analysis (min 48, max 340) when the two samples with low numbers were excluded (*S. betaceum* and *S. etuberosum*, Appendix S3). Furthermore, to reduce the effect of missing data and long branch attraction, sequences shorter than 25% of the average length for the gene were eliminated. The number of loci retained from the min04 and min20 datasets was 310 and 348 respectively, with the final aligned length varying between 242,272 bp and 261,975 bp (Appendix S10).

The effect of missing data was tested by comparing two different sampling thresholds based on the minimum number of taxa in each of the target genes alignments (min20 vs. min04, i.e. a minimum of 20 taxa per gene and a minimum of four taxa per gene, respectively) using HybPiper (Johnson et al., 2016) to retrieve and filter the genes.

ML analyses were run on both TC datasets in IQ-TREE2 (Minh et al., 2020) with partitioning between loci. In addition, IQ-TREE2 was used to generate individual ML trees for each loci, and the resulting phylogenetic trees were used for coalescent analyses with ASTRAL-III v.5.7.3 (Appendix S9; Zhang et al., 2018), where tree nodes with <10% BS values were collapsed using Newick Utilities v. 1.5.0 (Junier and Zdobnov, 2010). Trees with excessively long branches were identified using phyx (Brown et al., 2017) by looking at tree lengths and root-to-tip variation (command “pxlstr”); seven gene trees with excessively long branches were identified and excluded for the min20 and ten for the min04 datasets, leading to a total of 303 and 338 gene trees being used for the respective coalescent analyses. Branch support was assessed using local PP support (Sayyari and Mirarab, 2016) calculated in ASTRAL, where PP values >0.95 were considered strong, 0.75-0.94 weak to moderate, and ≤0.74 as unsupported.

### Discordance analyses

*Comparison of resulting species trees**–*** Topological congruence and discordances between all 10 topologies generated were assessed visually by generating graphical representations through custom R-scripts using the following packages: “ggtree” (Yu, 2020), “stringr” (Wickham and Wickham, 2019), “ape”(Paradis and Schliep, 2019), “ggplot2” (Villanueva and Chen, 2019) and “gridExtra”(Auguie and Antonov, 2017). To facilitate comparisons, all trees were reduced to include the outgroup *Jaltomata* and 9 taxa representing the following clades of *Solanum*, which were recovered in all analyses: Thelopodium, Regmandra, Potato, Morelloid (as a representative of both the Dulcamaroid and Morelloid clades), Archaesolanum, *S. anomalostemon* S.Knapp & M.Nee (species sister to Clade II), Acanthophora (minor clade of the Leptostemonum) and two representatives of the EHS clade (Table 1). The species sampled in the PL and TC datasets were identical for all except three minor clades, in which different closely related species were sequenced (Acanthophora: *S. viarum* Dunal/*S. capsicoides* All.; Morelloid: *S. opacum* A.Braun & C.D.Bouché/*S. americanum* Mill.)

*Concordance factors–* Phylogenomic discordance was measured using gene concordance factors (gCF) and site concordance factors (sCF) calculated in IQ-TREE 2 (Minh, Hahn, et al., 2020). These metrics assess the proportion of gene trees that are concordant with different nodes along the phylogenetic tree and the number of informative sites supporting alternative topologies. Low gCF values can result from either limited information (i.e., short branches) and/or genuine conflicting signal; low sCF values (∼30%) indicate lack of phylogenetic information in loci (Minh, Hahn, et al., 2020). The metrics were calculated using the TC ASTRAL min20 topology (303 genes) and the PL IQ-Tree topology of 151 species (unpartitioned) where sampling was reduced to 21 and 34 tips in TC and PL topologies, respectively, retaining a single tip for each of the different minor and major clades. An additional tip was retained for the Old Word Clade to visualize the gCF and sCF for the crown node of that lineage.

*Network analyses and polytomy tests–* The presence of reticulate evolution and conflicting signals in gene trees in the TC dataset was explored by generating a filtered supertree network in SplitsTree 4 (Huson and Bryant, 2006) of the TC min20 dataset (303 genes) collapsing branches with <75% local PP support with a minimum number of trees set to 50% (151 trees). Polytomy tests were carried out in ASTRAL-III (Sayyari and Mirarab, 2018) using ASTRAL topologies of the two datasets (min20 and min04). Gene trees were used to infer quartet frequencies for all branches to determine the presence of polytomies while accounting for ILS. The analysis was run twice to minimize gene tree error.

## RESULTS

### Phylogenetic analyses

*Congruent recovery of major clades–* All three datasets, including the supermatrix and the two phylogenomic datasets (PL and TC), recovered previously recognized major clades in *Solanum* (Fig. 1-2a,c); a few minor clades, concentrated in Clade II, were found to be polyphyletic in the supermatrix phylogeny, including the Mapiriense-Clandestinum, Sisymbriifolium, Wendlandii-Allophyllum and Cyphomandropsis minor clades (Appendices S11-S12); comparison with PL and TC phylogenies is not possible, as only one species of each clade were sampled in these datasets. In Clade I, nearly all specimens of the Dulcamaroid clade formed a monophyletic group. The only exception concerned *S. alphonsei* Dunal, sampled here for the first time. In both the supermatrix and PL analyses, this species was sister to *S. valdviviense of the* Valdiviense clade, with maximum branch support in the PL analyses (Fig 2., Appendix S13).

**Figure 1.**
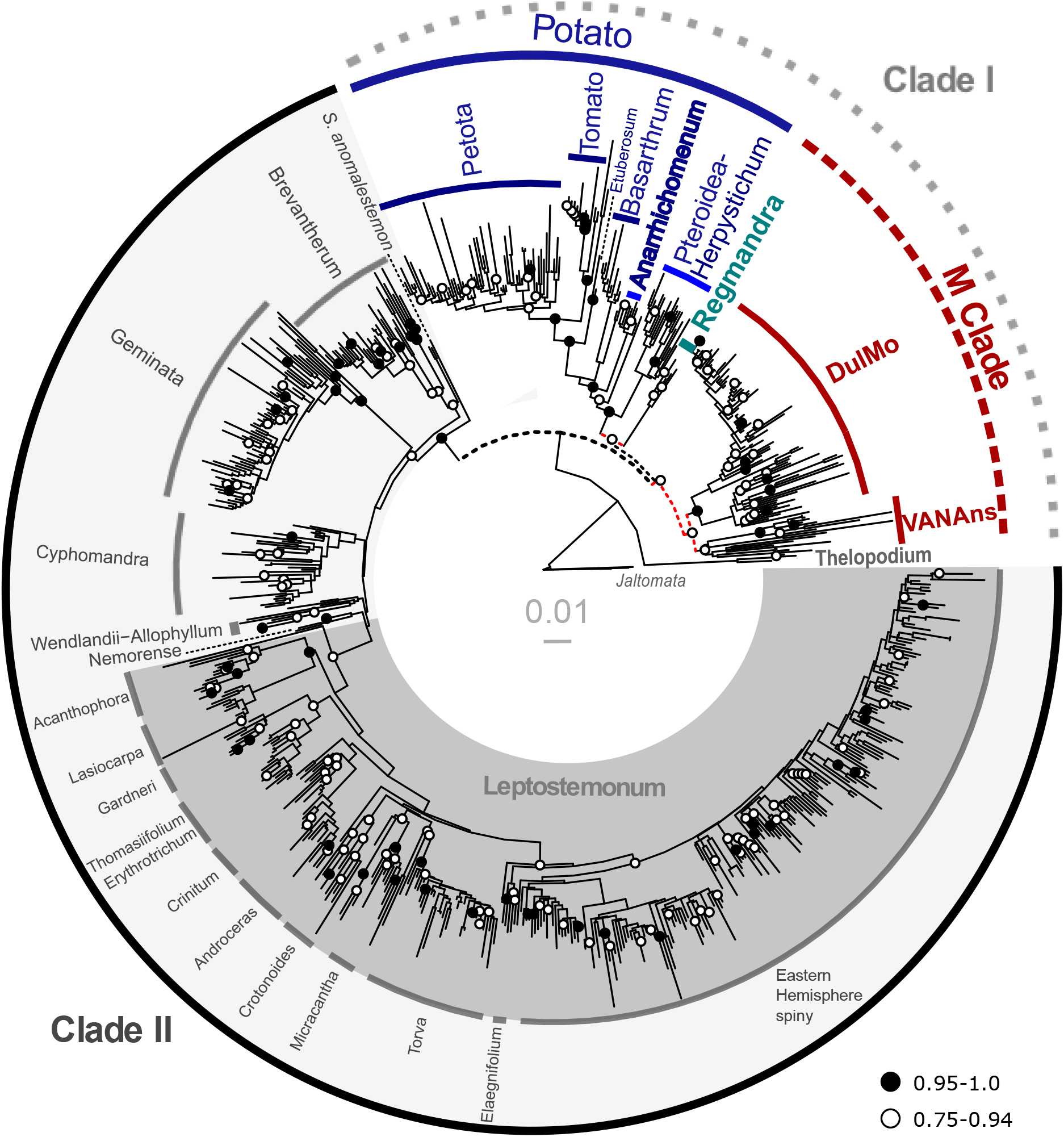
Supermatrix phylogeny from Maximum Likelihood analysis (RaxML) of 742 *Solanum* species based on two nuclear and seven plastid regions. Bootstrap branch support values are color coded: black = strong (0.95–1.0), white = moderate to weak support (0.75–0.94). Dashed lines indicate in phylogeny indicate relationships that were not recovered in the TC and PL analyses (see Figures 2-3). Clade names refer to major and minor clades discussed in the text (see Table 1); dashed lines for clade labels indicate groups that were not recovered in the TC and PL analyses.

**Figure 2.**
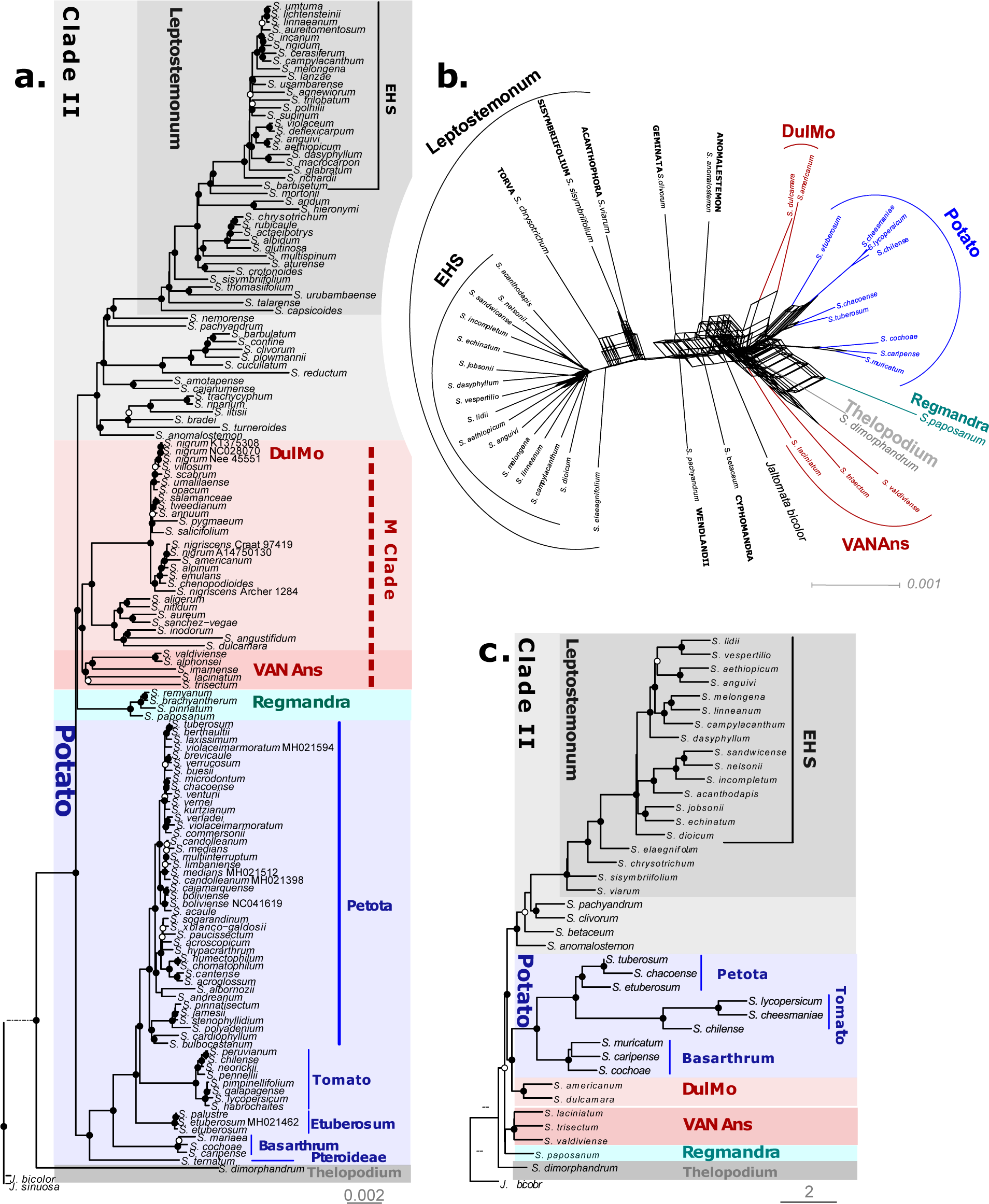
Comparison of *Solanum* clades recovered in plastome (PL) and target-capture (TC) phylogenomic datasets. a) Plastome phylogeny from the unpartitioned maximum likelihood analysis (PL-151-UP) based on 160 loci representing exons, introns and intergenic regions; b) Filtered supertree network of the TC dataset (min20) based on 303 gene trees with a 50% minimum tree threshold. c) TC phylogeny with 40 species from coalescent analysis (TC-min20-ASTRAL). Clades are shown in the same color in all three phylogenies to enable comparison. Branch support values (BS values in (a) and local PP values in (c)) are color coded: black = strong (0.95–1.0), white = moderate to weak (0.75–0.94). Scale bars = substitutions/site. Collection or Genbank numbers are indicated in the PL phylogeny for duplicate species sampled in the phylogenetic trees.

Despite these minor novelties, all analyses recovered the Thelopodium clade as sister to the rest of *Solanum* (Fig. 1-2, Appendices S11-15). The Potato clade was strongly supported across all analyses (Fig. 1-2, Appendices S11-S15), as was the Regmandra clade in supermatrix and PL analyses (only one sample in TC phylogenies). Furthermore, all analyses recovered a clade here referred to as DulMo that includes the Morelloid and Dulcamaroid clades (Figs. 1-2, Appendices S11-S15). A new strongly supported clade, here referred to as VANAns clade and comprising the Valdiviense (including *S. alphonsei*, see below), Archaesolanum, Normania, and the African non-spiny clades, was found across all analyses (Figs. 1-2; Appendices S11-S15).

Clade II was supported as monophyletic across all topologies (Fig. 1-2 a,c), with maximum branch support in all 10 species trees (Appendices S11-S15).While differences in sampling prevent thorough comparisons of relationships between clades within Clade II, there was no deep incongruences detected amongst topologies obtained with the supermatrix, PL, and TC datasets (Fig. 1-2 a,c; Appendices S9-S15). Within Clade II, the large Leptostemonum clade (the spiny solanums) was strongly supported in all cases (Fig. 1-2 a,c; Appendices S11-S15).

*Incongruent relationships amongst clades and impact of different analyses–* Overall, we found that despite using different phylogenetic analyses and investigating the impact of missing data and taxon sampling on the different datasets, these had little impact on the relationships recovered amongst clades. The BI and ML supermatrix analyses were identical in terms of composition and relationships of major clades (Fig. 3b), as were the four PL species trees (Fig. 3 d,e). There were some differences amongst the topologies of the TC datasets, but these differences concerned branches which had little support (Fig. 3a,c,b). Between supermatrix, PL and TC datasets, however, major incongruences between species trees were observed with respect to the relationships among the main clades identified in the section above (Fig. 1-3).

**Figure 3.**
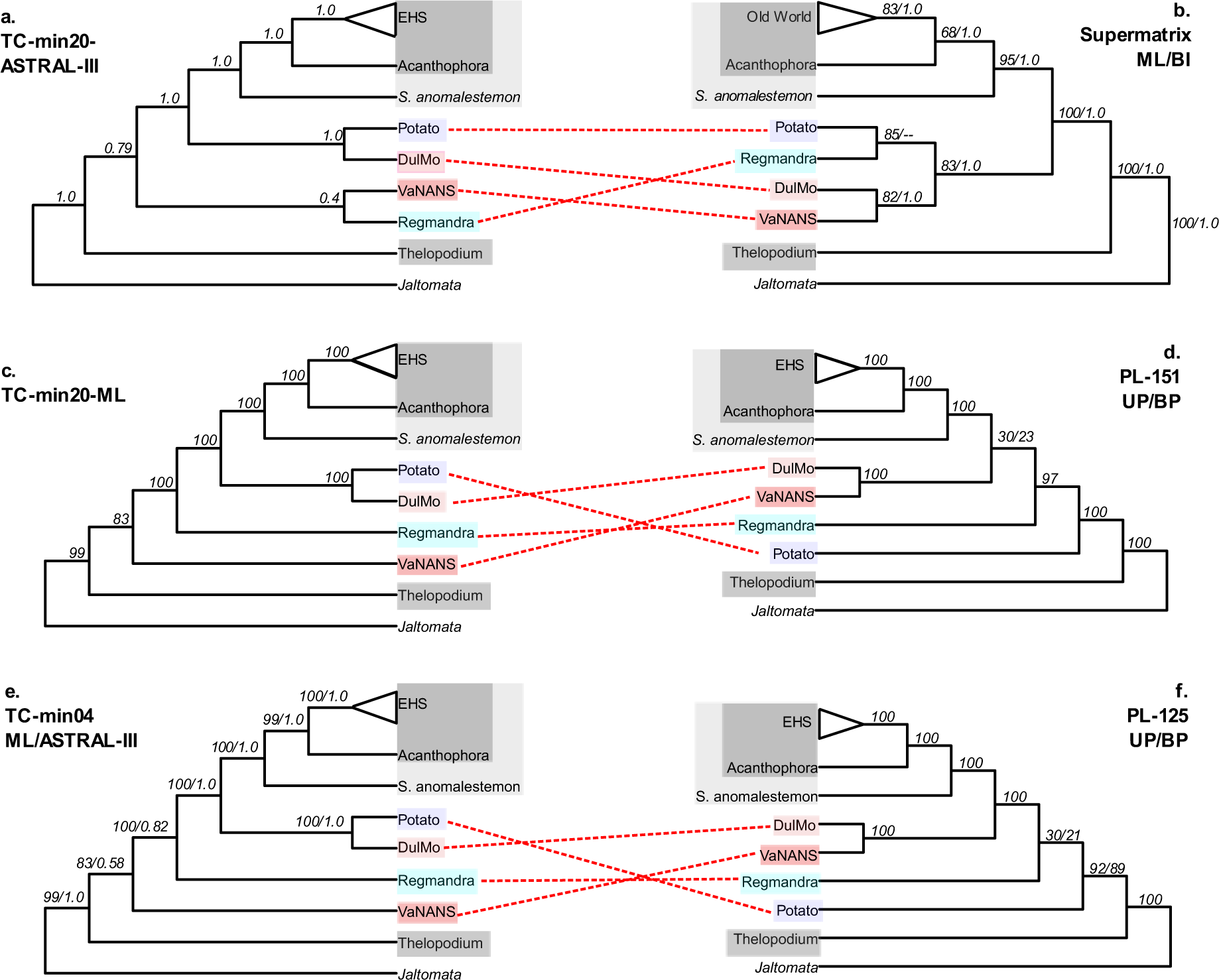
Comparison of *Solanum* clades recovered in the three different datasets. a) TC Astral-III phylogeny of the min-20 dataset, with local posterior probabilities indicated at nodes; b) ML and BI phylogenies of supermatrix dataset, with bootstrap support and posterior probabilities indicated at nodes; c) TC ML phylogeny of the min 20 dataset, with local posterior probabilities indicated at nodes; d) PL ML phylogenies of the unpartitioned and best partition- scheme of the 151 taxa dataset, with bootstrap for each respective analysis is indicated at nodes; e) TC ML phylogeny and Astral-III phylogeny of the min 04 dataset, with bootstrap support and local posterior probabilities indicated at nodes; f) PL ML phylogenies of the unpartitioned and best partition-scheme of the 125 taxa dataset, with bootstrap for each respective analysis indicated at nodes.

While the BI and ML supermatrix phylogeny supported the monophyly of the previously recognised Clade I that includes most non-spiny *Solanum* clades (Fig. 1, Appendices S11-12), the PL and TC phylogenetic trees resolved clades associated with Clade I as a grade relative to Clade II (Fig. 2a,c, Appendices S13-S15). This was in large part due to the unstable position of the Regmandra clade that was subtended by a particularly short branch and resolved in different positions along the backbone in all three datasets (Fig. 3). For example, the ML supermatrix analysis recovered the Regmandra clade as sister to the Potato clade with strong to moderate branch support (Fig. 3b), although the BI supermatrix analysis could not resolve whether the Regmandra clade was sister DulMo+VANAns clade or the Potato clade (Fig. 3b, Appendix S12). In contrast, the PL analyses resolved Regmandra as sister to the M clade + Clade II, with either maximal or no branch support at all (Fig. 3). The TC species trees resolved Regmandra as sister to the Potato clade, DulMo, and Clade II, with maximum support (Fig. 3). While one of the TC ASTRAL analysis also recovered this topology with moderate support (local posterior probability 0.82, Fig. 3), the other TC ASTRAL analysis resolved Regmandra as sister to the VANAns clade, but without any branch support (local PP 0.4, Fig. 3).

The previously identified M Clade composed of the VANAns and DulMo clades were not supported by all analyses (Fig. 3). While all PL ML analyses recovered the M clade with maximum BS values (Fig. 3), none of the TC analyses recovered it. Instead, they resolved the DulMo clade as sister to the Potato clade, with maximal BS or local PP support values (Fig. 3.) Furthermore, the VANAns clade was recovered as sister to the rest of *Solanum* (excluding the Thelopodium clade) with moderate support in the TC ML analyses. Placement of the VANAns clade in the TC-ASTRAL-III analyses had low or no support value, being resolved as either sister to DulMo, or sister to the rest of *Solanum,* excluding the Thelopodium clade (Fig. 3).

In addition, the position of the Potato clade within *Solanum* was incongruent between datasets: whereas it was resolved as sister to Regmandra in the supermatrix analysis, it was resolved as sister to the remaining *Solanum* in PL dataset, and sister to the DulMo clade in all TC analyses (Fig. 3), all with strong branch support. The phylogenomic datasets also showed incongruent positions for the Etuberosum clade within the larger Potato clade, where TC analyses resolved it as sister to the Petota clade with maximum local PP support in the ASTRAL analyses (Appendix S15); in the ML analyses, this position either had moderate BS values (76%) or was found to be nested within the Petota clade with no branch support (Appendix S14). In contrast, PL analyses placed Etuberosum clade as sister to the Tomato clade with maximum branch support (Appendix S13).

Finally, the BI and ML supermatrix phylogenies resolved the morphologically unusual *S. anomalostemon* as sister to the rest of Clade II (BS 95%, PP 1.0; Fig. 3, Appendices S11-S12). This contrasts with results from previous analyses, which found it to be part of the Mapiriense clade (Särkinen et al., 2015). PL analyses supported *S. anomalostemon +* Brevantherum clade as sister to the rest of Clade II with high branch support (Appendix S13). *Solanum anomalostemon* was also found to be sister to Clade II, although the Brevantherum clade was not included in the TC analyses preventing a strict comparison (Fig. 3). Two other taxa were found to represent single species lineage: *S. polygamum* as sister to the Leptostemonum clade and *S. euacanthum* as sister to the EHS clade (Appendices S11-S12). Within the Leptostemonum clade, the EHS clade was strongly supported in all analyses (Figs.1-3). There were however some minor differences in species-level relationships for closely related species of the Eggplant clade and Anguivi Grade (viz. *S. campylacanthum* Hochst. ex A.Rich., *S. melongena* L., *S. linnaeanum* Hepper & P.-M.LJaeger, *S. dasyphyllum* Schum. & Thonn. and *S. aethiopicum* L.; Fig. 1-2a, c, Appendices S11-S15).

### Discordance analyses

*Concordance factors**–*** Phylogenomic discordance was generally high across the PL and TC topologies, with gCF values >50% in only three nodes in the PL phylogeny (*Solanum* as a whole, *S. chilense* (Dunal) Reiche + *S. lycopersicum* L. or the Tomato clade, and *S. hieronymi* Kuntze + *S. aridum* Morong in the Leptostemonum clade; Fig. 4). Elsewhere, along the backbone of the PL phylogeny, gCF fell to 39% and below (8 nodes with gCF values 10% and below), with the lowest values found near branch nodes that varied the most amongst the different reconstructed species trees. This included the node subtending Regmandra (gCF 4%, SCF 38%; Fig. 4), and that positioning Regmandra + DulMo + VANAns clade as sister to Clade II (gCF 2%, SCF 31%). Similarly, low gCF and uninformative sCF values around 33% were found across Clade II, including the node placing *S. hieronymi* + *S. aridum* as sister to the Elaeagnifolium + Old World Minor clades (gCF 6 %, sCF 36 %; Fig. 4), as well as the placement of the Erythrotrichum + Thomasiifolium clades within the large Leptostemonum clade (gCF 5%, sCF 23%; Fig. 4).

**Figure 4.**
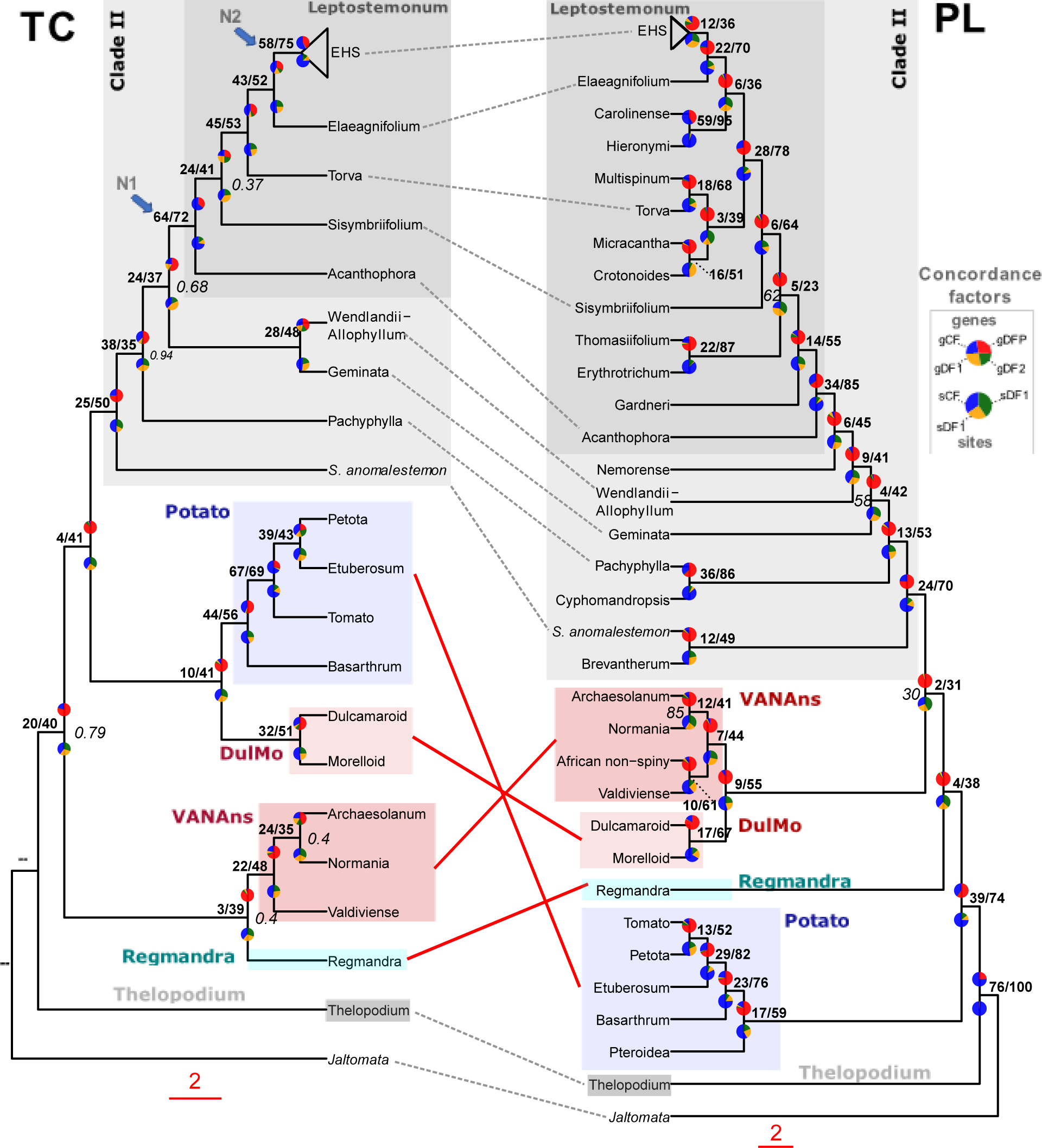
Discordance analyses within and between the plastome (PL) and target capture (TC) phylogenomic datasets across *Solanum*. Rooted TC ASTRAL phylogeny (left) and PL IQ-TREE2 phylogeny (right) with gene concordance factor (gCF) and site concordance factor (sCF) values shown as pie charts, above and below each node respectively; the PL topology is the unpartitioned ML analysis of 151 taxa, whereas the TC topology is based on the analysis of 40 taxa and 303 genes recovered from the A353 bait set. Both trees have been pruned to retain a single tip for each of the major and minor clades present within the PL and TC datasets. For gCF pie charts, blue represents proportion of gene trees concordant with that branch (gCF), green is proportion of gene trees concordant for 1^st^ alternative quartet topology (gDF1), yellow support for 2^nd^ alternative quartet topology (gDF2), and red is the gene discordance support due to polylphyly (gDFP). For the sCF pie charts: blue represents proportion of concordance across sites (sCF), green support for 1^st^ alternative topology (quartet 1), and yellow support for 2^nd^ alternative topology (quartet 2) as averaged over 100 sites. Percentages of gCF and sCF are given above branches, in bold. Branch support (local posterior probability) values ≥0.95 are not shown, and 0.94 and below are shown in italic grey, on the right; double-dash (--) indicates that the branch support was unavailable due to rooting of the phylogenetic tree. The nodes corresponding to the two clear clades identified in the filtered supertree network (see Fig. 2b) are identified as “N1” and “N2”.

Across the TC phylogeny, gCF and sCF values were slightly higher on average, with 3 nodes presenting values >50% for both metrics: one within the Petota clade (gCF 67%, SCF 69%; Fig. 4), one at the base of the Leptostemonum clade (gCF 64%, SCF 72%; Fig. 4), and another at the base of the EHS clade within Leptostemonum (gCF 58%, SCF 75%; Fig. 4). Three nodes had low gCF values of 10% or less, with again some of the lowest values located near the base of the tree, including the relationship of Regmandra as sister to the VANAns clade (gCF 3%, sCF 39%; Fig. 4), or placement of Potato as sister to the DulMo clade (gCF 10%, sCF 41%; Fig. 4), and the relationship of the Potato + DulMo clades as sister to Clade II (gCF 4%, sCF 41%; Fig. 4).

*Network analyses and polytomy tests–* High amount of reticulation/gene tree conflict was recovered between major clades of *Solanum* previously assigned to Clade I (e.g., Thelopodium, Regmandra, Potato, DulMo, VANAns), as well as with some lineage belonging to Clade II in the filtered supertree network using the TC data with 303 genes (min20; Fig. 2b). The network clearly supported the monophyly of the Leptostemonum and the EHS clade (Fig. 2b), corresponding to the nodes with high gCF and sCF values in the TC ASTRAL phylogeny (N1 and N2, Fig. 4).

The polytomy tests carried out for the two TC ASTRAL datasets resulted in 10 nodes each for which the null hypothesis of branch lengths equal to zero was accepted, suggesting they should be collapsed into polytomies (Appendix S16); these nodes corresponded to the ones subtending the Regmandra, Leptostemonum and EHS clades, but were also located within the VANAns clade as well as within Clade II, the. Polytomies were also detected with the Petota clade, including at the base of the Tomato clade (min4 dataset, Appendix S16), and at the base of the Etuberosum + Petota +Tomato clade (min20 dataset, Appendix S16). Repeating the analysis by collapsing nodes with <75% local PP support led to the collapse of 12 to 13 nodes across the analyses, most of them affecting the same clades as in the previous runs, but also leading to the collapse of the crown node of *Solanum*. The effective number of gene trees was too low when nodes with <75% local PP support were collapsed to carry out the test for two nodes subtending *S. betaceum* and *S. anomalostemon*, most likely related to the low number of genes recovered for *S. betaceum* (Appendix S3).

## DISCUSSION

The results of the ten phylogenetic analyses conducted here provide an updated evolutionary framework for the large and economically important genus *Solanum,* demonstrating that the major and minor clades within the group are stable (with a few noteworthy exceptions, see below). However, the strong levels of nuclear and nuclear-plastome discordance uncovered in the PL and TC analyses, in combination with the network analysis and polytomy tests, suggest that there are polytomies present along the backbone of the phylogeny. We first discuss the stability of the clades within *Solanum*, and the discovery of a few novel minor clades. We then examine the nuclear-plastome discordance and polytomies recovered and explore the possible causes underlying these, and their implications for the study of biogeography and trait evolution.

### Updated evolutionary framework for *Solanum*

The supermatrix phylogeny, despite being based on only nine loci, nearly doubles the species sampling, confirming the monophyly of most major and minor clades established in previous analyses (Särkinen et al. 2013) and the polyphyly of three minor clades (Pachyphylla, Cyphomandropsis, and Allophyllum, the latter including species of Mapiriense-Clandestinum clade). It also reveals three new minor clades in *Solanum* comprising a single species each and confirms the placement of 129 previously unsampled species (e.g., *S. alphonsei* in the Valdiviense clade and *S. graveolens* in the Cyphomandra clade; Appendices S11-S12).

Meanwhile, the phylogenomic analyses with increased gene sampling reveal a previously undetected major clade referred to as VANAns comprising of four minor clades (Valdiviense, Archaesolanum, Normania, and African non-spiny clades). Finally, our results did not support two previously resolved major clades due to nuclear-plastome discordance (Clade I and the M clade; Fig. 2). Detailed molecular systematic studies with increased taxon and genetic sampling will be required to fully resolve the circumscription of all the major and minor clades recovered with diagnostic features, including the new ones identified here (Hilgenhof et al. in prep).

Overall, our results establish that the taxonomic framework used in *Solanum* dividing the large genus into major and minor clades is robust, based on both phylogenomic datasets recovering the same major clades independent of methodological choices compared to the Sanger sequence supermatrix (e.g., Thelopodium, Regmandra, Potato, DulMo, VANAns, Clade II, Leptostemonum, and EHS clade). The major and minor clades currently used as informal infrageneric groups in *Solanum* were first established by Bohs (2005) based on a single locus of c. 2,000 bp in length (*ndhF*). Our results demonstrate that larger species and gene sampling support the clades established nearly two decades ago (e.g. Weese and Bohs, 2007; Särkinen et al., 2013). However, increased gene sampling provided by the two phylogenomic datasets does not help to resolve any of the polytomies along the backbone of *Solanum* close to the crown node and along the backbone of Clade II (Särkinen et al., 2013).

### Nuclear & nuclear-plastome discordance

Our results reveal three regions of the *Solanum* phylogeny with gene discordance with low gCF and sCF values in the PL and TC dataset (Fig. 4). These regions with nuclear discordance include: (1) the backbone of *Solanum* near the crown node of the genus where major clades previously identified as Clade I diverge (from here on referred to as Grade I); (2) the backbone of the large Leptostemonum clade; and (3) the backbone of the EHS clade within the Leptostemonum (Figs. 2b and 3). Many of the branches within these regions are extremely short in both PL and TC phylogenomic datasets (Fig. 1-2, Appendices S11-S15), and network analyses of the nuclear dataset reveals reticulation in one of them (Grade I, Fig. 2b). Polytomy tests confirm that multiple nodes within all three regions should be collapsed in the TC dataset (Appendix S16) and support the recognition of these regions as polytomies. Hence, we refer to these three regions of the phylogeny as polytomies from hereon.

Further exploration of the polytomies reveal nuclear-plastome discordance within Grade I, relating to the position and relationship between Regmandra, Potato, DulMo and VANAns clades (Fig. 3-4). No signal of nuclear-plastome discordance was detected in the other polytomies based on the species sampling presented here (Fig. 3-4), but increased species sampling will be needed to confirm these results.

Altogether, our results indicate the presence of three polytomies which differ somewhat in nature. The deepest of these polytomies along the backbone of *Solanum* near the crown node shows high nuclear and nuclear-plastid discordance with reticulation evident even within the nuclear phylogenomic dataset (Fig. 2b). This polytomy could be referred to as a hard polytomy, because it will probably be difficult to resolve even with more genomic data, due to its deeper position in the phylogeny in terms of evolutionary depth and time, the presence of clear nuclear-plastome discordance, short branch lengths and evidence for reticulation within the nuclear phylogenomic dataset. In contrast, the other two polytomies along the backbone of Leptostemonum and the EHS clades are at shallower evolutionary depth and show nuclear discordance only without clear/widespread reticulation in the nuclear dataset (Fig. 2b). These polytomies represent simpler cases and may turn out to be possible to resolve with more genomic data. In either case, to confirm whether the polytomies recovered here are truly “hard” or “soft”, denser taxon sampling and more genomic data will be required to carry out more rigorous tests concerning the cause of the gene discordance observed here.

### What is causing genomic discordance in our dataset?

Finding genomic discordance in our phylogenomic datasets is unsurprising, given that it has also been found in many other phylogenomic studies in the Solanaceae, including *Nicotiana* (Dodsworth et al., 2020), the Capsiceae (*Capsicum* and relatives; Spalink et al., 2018), subtribe Iochrominae (Gates et al., 2018), *Jaltomata* (Wu et al., 2019), and two studies of *Solanum* involving the Tomato (Strickler et al., 2015; Pease et al., 2016) and Petota clades (Huang et al., 2019). ILS was shown to be responsible for the widespread discordance found in phylogenomic data in the diploid Tomato clade (Strickler et al., 2015; Pease et al., 2016), while hybridization and introgression has been argued to be behind genomic discordance in Petota clade that includes many polyploids (Huang et al., 2019).

Potential processes responsible for nuclear or nuclear-plastome discordance involve gene introgression, ILS, hybridization, and polyploidisation; distinguishing between these remains difficult even with increased genomic sampling involving custom bait sets (Larridon et al., 2020; Koenen et al., 2021) or whole genome-sequences (Suh, 2016; Malinsky et al., 2018; Williams et al., 2021). Comparison of the nuclear and plastome topologies in our study does not indicate any obvious chloroplast capture events that could explain the observed nuclear-plastome discordance along the backbone of *Solanum* near the crown node. Furthermore, cytogenetic and chromosome studies show no evidence for genome duplication or polyploidy along the three polytomies discovered here, despite the three-fold increase in genome size between the distantly related potato (*S. tuberosum* L., Potato clade) and eggplant (*S. melongena,* Leptostemonum clade; Barchi et al., 2019). Chromosome counts indicate that the ancestor of *Solanum* was diploid: a large majority of *Solanum* species are reported to be diploid (>97% of the 506 species for which chromosome counts are available), and mapping of ploidy level across the phylogeny indicates that most of the lineages involved in the three polytomy regions identified here are diploid (Chiarini et al., 2018). Polyploidy has arisen independently within the Archaesolanum, Petota, Morelloid, Caroliniense, Elaeagnifolium and EHS minor clades within the larger Leptostemonum clade (Chiarini et al., 2018), and hybridization/introgression has been argued to be the case behind phylogenomic discordance found in the Petota clade (Huang et al., 2019).

Gene duplication could explain the signal recovered here for the EHS clade but is unlikely to explain the discordance observed here. Save for one locus, our analyses did not detect the presence of paralogs in our nuclear dataset.

Currently, the most likely explanation for the discordance along the backbone of *Solanum* is due to ILS caused by rapid speciation. Two of the polytomies include the most species-rich (Table 1) and rapidly diversifying lineages of *Solanum*, the Leptostemonum and the EHS clades (Echeverría-Londoño et al., 2020), whose crown ages have been estimated to be between 8-11 and 4-6 million years (Myr), respectively (Särkinen et al., 2013). The backbone of *Solanum* near the crown node has been estimated to be almost twice as old as the Leptostemonum clade (13-17 Myr; Särkinen et al. 2013) yet shows a strong signal of nuclear-plastome discordance. While past studies have not detected any increased rates of diversification near the crown node of *Solanum*, detecting diversification rate shifts remains a challenge (Louca and Pennell, 2020), especially in older nodes. Hence, we cannot fully exclude the option that ILS and rapid speciation has taken place close to the crown node of the genus.

Presence of short internal branches is typical of ILS in lineages with large population sizes and high mutation rates (Schrempf and Szöllősi, 2020). This fits with the biology of *Solanum* in general, which is typically known to contain “weedy”, disturbance-loving pioneer species resilient to change. Many species are known to have large geographical ranges and ecological amplitude, including globally distributed weeds from the Leptostemonum, Brevantherun and Morelloid clades, such as *S. elaeagnifolium, S. caroliniense*, *S. torvum*, *S. erianthum, S. mauritianum, S. americanum* and *S. nigrum* (Knapp et al., 2017, 2019; Cowie et al., 2018; Särkinen et al., 2018). Some of the weedy characteristics found in these species include the ability to improve fitness and defense traits in response to disturbance (Chavana et al., 2021), as well as having allelopathic properties which allow them to establish themselves to the detriment of native vegetation (Cowie et al., 2018). If such characteristics were present in ancestral *Solanum*, they could have promoted rapid speciation across the globe, followed by rapid morphological evolution and speciation within areas. The patterns observed here could possibly be the result of three major rapid speciation “pulses” across the evolutionary history of *Solanum,* involving lineages close to the crown node of *Solanum*, Leptostemonum, and the EHS clade. The idea of an ecologically opportunistic ancestor is supported by the tendency of many of the major clades near the crown node of *Solanum* to occupy periodically highly stressed and disturbed habitats, including flooded varzea forests occupied by Thelopodium clade, hyper-arid deserts occupied by Regmandra clade, and highly disturbed and dynamic open mid-elevation Andean montane habitats occupied by DulMo clade, where landslides are among the most common areas where many of the species are found (Knapp, 2013; Särkinen et al., 2018; Knapp et al., 2019).

Future studies with larger datasets will be able to carry out additional tests, such as the impact of using phylogenetic models that take into consideration the heterogeneity of molecular sequence evolution (Williams et al., 2021), as well as different data types (Romiguier et al., 2013; Reddy et al., 2017). Future studies will need to untangle how introgression and ILS are potentially affecting the patterns of genomic discordance observed here at different phylogenetic depths (Meleshko et al., 2021). Additional information about recombination, chromosome structure, and genomic size and evolution of *Solanum* will also be useful to clearly define coalescence genes in phylogenomic datasets, fundamental units in coalescent analyses which are rarely examined (Springer and Gatesy, 2018). Currently, information about genome evolution in *Solanum* is lacking, as only 62 species (5% of *Solanum*) are recorded in the plant DNA C-value database (Pellicer and Leitch, 2020), and 86 species (7% of *Solanum)* have been studied with chromosome banding and/or FISH techniques (Chiarini et al., 2018). Information about genome size is missing for lineages such as the Thelopodium and Regmandra clades and for the majority of species not directly related to major commercial crops.

### Implications for biogeographical and morphological studies in Solanum

The idea that well-supported and fully bifurcating phylogenies are a requisite for evolutionary studies is built on the premise that such trees are the accurate way of representing evolution. The shift in systematics from “tree”- to “bush”-like thinking, where polytomies and reticulate patterns of evolution are considered as acceptable or real (Poczai, 2013; Mallet et al., 2016; Edelman et al., 2019), comes from the accumulation of studies finding similar unresolvable phylogenetic nodes, despite using different large-scale genomic sampling strategies and various analytical methods (Suh, 2016). Given the difficulty of resolving short internal branches in phylogenies and the rapid evolution of major clades in *Solanum,* it will be important to adopt methods that incorporate polytomies and networks to conduct biogeographical and morphological studies (Than et al., 2008; Solís-Lemus et al., 2017; Wen et al., 2018; Olave and Meyer, 2020; Lutteropp et al., 2021).

In terms of biogeography, our inability to resolve relationships amongst the major lineages in *Solanum*, especially along the backbone of *Solanum* near the crown node, has implications for understanding the ancestral environment of *Solanum* and its major lineages. Uncertainty amongst the relationships of major clades does not change the hypothesis that the genus probably originated from South America and spread multiple times to Africa, Asia, Australia, North America, and Europe (Olmstead and Palmer, 1997; Echeverría-Londoño et al., 2020). The polytomy near the crown node of *Solanum* does, however, cast uncertainty on the specific region and habitat/biome that the major clades originated within the South American continent. For example, the sister relationship of Regmandra and the Potato clade inferred by the Sanger supermatrix analysis suggests that the wild ancestors of both potato and tomato evolved from an ancestor adapted to survive in lomas deserts from coastal South America (Bennett, 2008; Fig. 1). Yet, both nuclear and plastome phylogenomic datasets suggest that the Potato clade is more closely related to the DulMo clade found to occur in tropical montane and subtropical biomes (Fig. 3)

The hard polytomy along the backbone of *Solanum* also has important implications for evolutionary biologists interested in trait evolution. Standard methods of trait evolution relying on bifurcating trees may incorrectly infer how traits evolve (Hahn and Nakhleh, 2016). The discordance between traits, gene trees, and species trees has been defined as hemiplasy (Avise and Robinson, 2008), and studies have shown that depending on the level of ILS present in the data, hemiplasy can lead to different interpretations of convergent evolution of traits across phylogenetic trees (Mendes et al., 2016). While broad mapping of morphological traits on a species-level phylogeny can help gain a rough understanding of phenotypic variation across clades, careful study of gene tree topologies in relation to a trait of interest is essential to gain an exact understanding of its evolutionary origin.

Our findings reflect results from recently published studies showing rapid morphological innovation coinciding with areas of strong phylogenomic discordance in different plants and animal groups (Parins-Fukuchi et al., 2021), where the signal of nuclear-plastome discordance corresponds to strong ecological diversification and morphological innovation across major clades in *Solanum* previously assigned to Clade I. The major clades involved in the nuclear-plastome discordance along Grade I show large differences in their ecology as well as morphology. Members of the Thelopodium, Regmandra, VANAns, Potato, and DulMo clades occupy a wide range of tropical, montane and temperate habitats across South America, Africa and Australia (Symon, 1994; Knapp, 2000; Bohs and Olmstead, 2001; Spooner et al., 2004, 2016, 2019; Bohs, 2005; Peralta et al., 2007; Bennett, 2008; Knapp, 2013; Knapp and Vorontsova, 2016; Tepe et al., 2016; Särkinen et al., 2018; Knapp et al., 2019). Morphology shows equally high polymorphism between these major clades across many traits, such as growth form, which varies from single-stemmed wand-like shrubs (Thelopodium clade), annual herbs (Regmandra, Potato, and Morelloid clade), woody climbers and shrubs (VANAns clade), and herbaceous vines rooting along nodes (Potato clade). Similar patterns are observed in inflorescence position and branching, corolla shape, stamen dimorphism, and anther shape showing the presence of high polymorphism in these clades of which only some was retained in Clade II (Hilgenhof et al. in prep). Testing the idea that this phenotypic diversity is linked to ecological diversification will require the construction of detailed morphological and ecological datasets to test if this pattern holds up in more formal and rigorous analyses.

## CONCLUSION

We demonstrate the stability of the majority of the clades defined within *Solanum* and uncover significant nuclear and nuclear-plastome discordance amongst relationships of major clades in *Solanum* based on the first phylogenomic study of the genus with wide species sampling. Three major polytomies are identified in *Solanum* based on the short branch lengths, gene concordance factor results, and polytomy tests. Two of these polytomies correspond to the biggest and most quickly diversifying lineages within *Solanum* (Leptostemonum and EHS clades). The third polytomy along the backbone of *Solanum* near the crown node involves reticulation and strong nuclear-plastome discordance and highlights great uncertainty in the relationships between the Potato, DulMo, Regmandra, and VANAns clades. This region of nuclear-plastome discordance corresponds with high ecological and morphological innovation and we argue that it is most likely due to ILS and rapid speciation based on current knowledge of genome evolution in *Solanum*. Future studies, even with full genome sequences and increased taxon sampling, might not be able to resolve the polytomy near the crown node of *Solanum* because the pattern of high reticulation combined with internodal short branches and it’s older age. Data on genome size and chromosome structure of the earliest branching lineages in *Solanum* will be required to further explore the nature and causes of this hard polytomy. We argue that acknowledging and embracing polytomies and reticulation is crucial if we are to design research programs aimed at understanding the biology of large and rapidly radiating lineages, such as the large and economically important *Solanum*.

## Acknowledgements and funding

This work was supported by the Fonds de recherche du Québec en Nature et Technologies postdoctoral fellowship and a grant from the Department of Biological Sciences of the University of Moncton to EG, the Sibbald Trust fellowship to RH, the Ceiba Foundation to AO, CNPq Conselho Nacional de Desenvolvimento Científico e Tecnológico awards 479921/2010-54 and 427198/2016-0 and Coordenação de Aperfeiçoamento de Pessoal de Nível Superior CAPES/FAPESPA award 88881.159124/2017-01 to LLG, NSF through grant DEB- 0316614 “PBI Solanum: a worldwide treatment” to SK and LB, the Calleva Foundation & Sackler Trust (Plant and Fungal Trees of Life Project at Kew), the LUOMUS Trigger and Systematics Research Fund to PP, the OECD CRP and Eötvös Research Grant (MAEÖ-00074- 002/2021). Field sampling was supported by the Northern Territory Herbarium and the David Burpee Endowment at Bucknell University (Australia), and National Geographic Society Northern Europe Award GEFNE49-12 (Peru, TS). Peruvian specimens were collected and sequenced under the permission of Ministerio de Agricultura, Dirección General Forestal y de Fauna Silvestre (collection permits 084-2012-AG-DGFFSDGEFFS and 096-2017- SERFOR/DGGSPFFS, and genetic resource permit 008-2014-MINAGRI-DGFFS/DGEFFS).

We thank Elliot Gardner for sharing scripts and advice on phylogenomic analyses with HybPiper, Royce Steeves for providing advice on DNA extraction for genome skimming, Felix Forest and Olivier Maurin for providing technical support and providing feedback on the manuscript, and João R. Stehmann, Thais Almeida, Paul Gonzáles, and Maria Baden who greatly contributed to fieldwork and sample acquisition. Finally, we would also like to thank the three reviewers, including Stacey Smith and William J. Baker, who provided constructive reviews and feedback that greatly improved the final version of this manuscript.

## Author contributions

EG designed and performed the analyses of the paper, with guidance from PP, AO, SD and TS; EG produced all figures, and wrote the article, with major contributions from TS, and PP, SD, SK and XA. RH and TS helped in data gathering and analyses. All other authors contributed data to the main analyses. All authors read and contributed to the final version of the manuscript.

## Data Availability Statement

Raw sequence data generated in this study are deposited in various archives, including Genbank (https://www.ncbi.nlm.nih.gov/genbank/) and the European Nucleotide Archive (https://www.ebi.ac.uk/ena/browser/home); full accession numbers are provided in Appendices S1, S2 and S3. In addition, the 10 species trees generated for this study, as well as the alignments used for the different phylogenetic analyses, including the concatenated Sanger supermatrix, the plastome dataset and and the target capture datasets (min 04 and min20) are available via Data Dryad, at the following link: (to be provided upon acceptance for publication).

## Supporting Information

Additional supporting information may be found online in the Supporting Information section at the end of the article:

Appendix S1. Supermatrix sample information, including voucher details and Genbank numbers for sequences used.

Appendix S2. Plastome (PL) sample information, including voucher details and plastome assemblies’ results. Total length, as well as length for the long-single copy region (LSC), the short-single copy region (SSC) and the two inverted repeat regions (IR1 and IR2) is shown; statistics of mean coverage per base pair and standard deviation are also provided.

Appendix S3. Target-capture (TC) sample information, including voucher details and sequence recovery statistics. The number of reads (NumReads), the number of reads mapped to the targets

(ReadsMapped), the percentage of reads on target (PctOnTarget), the number of genes with reads (GenesMapped), the number of genes with contigs (GenesWithContigs), (GenesWithSeqs, GenesAt25pct, GenesAt50pct, GenesAt75pct, GenesAt150pct, and the number of genes with paralog warnings (ParalogWarnings) is shown.

Appendix S4. Supermatrix alignment details, with details about the nine regions selected for this study. Number of species sampled per region, accumulative percentage of species sampled per region, aligned length, proportions of parsimony informative characters (PI), and variable sites (VS) per region in the dataset are indicated. Values are calculated with outgroups, and with ambiguous regions and repeats excluded. bp=base pairs.

Appendix S5. List of polyploid taxa in *Solanum*.

Appendix S6. ML results for each of the nine individual loci and combined plastid loci. (a) ITS; (b) matK; (c) ndhF; (d) ndhF-rpL32; (e) psbA-trnH; (f) rpL32-trnL; (g) trnL-trnT; (h) trnS-trnG; (i) waxy; (j) seven plastid loci. Nodes with bootstrap support equal and above 95% are in cyan, and with branch support between 75% and 94% in red. Tips indicate species names, followed by major and/or minor clade, as indicated in Table 1.

Appendix S7. BI results for each of the nine individual loci and combined plastid loci. (A) ITS; (B) matK; (C) ndhF, (D) ndhF-rpL32, (E) psbA-trnH (F) rpL32-trnL; (G) trnL-trnT; (H) trnS-trnG; (I) waxy; (J) seven plastid loci. Nodes with posterior probability equal and above 0.95 are in cyan, and nodes with posterior probabilities between 0.75 and 0.95 are in red. Tips indicate species names, followed by major and/or minor clade, as indicated in Table 1.

Appendix S8. Plastome (PL) alignment statistics for plastome alignment. Data shows number of sequences, trimming mode, the number of loci retained for coalescent analysis after checking for excessive gene tree branch lengths, alignment length, number of informative and constant sites, pairwise identity, average GC content, percentage of gaps, and average locus length for the exon, intron and intergenic regions.

Appendix S9. Optimal substitution model used in ML analyses for the PL and TC datasets, determined using ModelFinder in IQ-TREE2. For each locus, the number of taxa, sites, informative sites, and invariable sites are indicated, as well as the model selected and the AICc score. Worksheet titles correspond to the following: PLUnpartitioned = Models selected for PL unpartitioned datasets, for 151 taxa and 125 taxa; PLBestPartScheme = Models selected for PL datasets analysed according to the best-partition scheme; TCPartitioned_Min4= Models selected for loci of the TC dataset, with minimum 4 taxa per loci; TCPartitioned_Min20= Models selected for loci of the TC dataset, with minimum 20 taxa per loci.

Appendix S10. Target-capture (TC) alignment statistics. Loci excluded refer to the number of excluded loci based on excessively long branch lengths, and loci retained is the final number of loci retained for both ML and coalescent analyses. Empty sequences inserted refers to amount of missing data. Min = minimum; Bp = base pairs. Appendix S11. Detailed RaxML of supermatrix phylogenetic tree with 746 taxa. Nodes with bootstrap support equal and above 95% are in cyan, and with branch support between 75% and 94% in red. Bootstrap support values for each node indicated in italic. Tips indicate species names, followed by major and/or minor clade, as indicated in Table 1.

Appendix S12. Detailed Bayesian inference (Beast) supermatrix phylogenetic tree with 746 taxa. Nodes with posterior probability equal and above 0.95 are in cyan, and nodes with posterior probabilities between 0.75 and 0.95 are in red. Posterior probability values for each indicated in italic. Tips indicate species names, followed by major and/or minor clade, as indicated in Table 1.

Appendix S13. ML phylogenetic trees of plastome datasets. Nodes with bootstrap support equal and above 95% are in cyan, and with branch support between 75% and 94% in red. Tips indicate species names, followed by major and/or minor clade, as indicated in Table 1 a) 151 taxa, all data, unpartitioned; b) 125 taxa, all data, unpartitioned; c) 151 taxa, all data, best partition scheme; d) 125 taxa, all data, best partition scheme.

Appendix S14. ML phylogenetic trees of A353 target capture datasets (IQ-TREE2). Nodes with bootstrap support equal and above 95% are in cyan, and with branch support between 75% and 94% in red. Tips indicate species names, followed by major and/or minor clade, as indicated in Table 1. a) filtering threshold of a minimum of 4 taxa per loci; b) filtering threshold of a minimum of 20 taxa per loci.

Appendix S15. Coalescent phylogenetic trees of A353 target-capture datasets (ASTRAL-III). Nodes with multi-locus local posterior probability support equal and above 0.95 are in cyan, and with support between 0.75 and 0.94 in red. Tips indicate species names, followed by major and/or minor clade, as indicated in Table 1. a) filtering threshold of minimum of 4 taxa per loci;) filtering threshold of minimum of 20 taxa per loci.

Appendix S16. Polytomy test results with ASTRAL-III. a) Target Capture A353 species tree ASTRAL-III, filtering threshold of minimum 4 taxa per loci, branches in gene trees with 10% or less branch support collapsed; b) Target Capture A353, ASTRAL-III, filtering threshold of minimum 4 taxa per loci, branches in gene trees with 75% or less branch support collapsed; c) Target Capture A353, ASTRAL-III, filtering threshold of minimum 20 taxa per loci, branches in gene trees with 10% or less branch support collapsed; d) Target Capture A353, ASTRAL-III, filtering threshold of minimum 20 taxa per loci, branches in gene trees with 75% or less branch support collapsed;

## References

Aberer, A. J., D. Krompass, and A. Stamatakis. 2013. Pruning rogue taxa improves phylogenetic accuracy: An efficient algorithm and webservice. Systematic biology 62: 162–166.

Amiryousefi, A., J. Hyvönen, and P. Poczai. 2018. The chloroplast genome sequence of bittersweet (Solanum dulcamara): Plastid genome structure evolution in Solanaceae. PloS one 13: e0196069.

Andrews, S., and Others. 2010. FastQC: a quality control tool for high throughput sequence data. APG. 1998. An ordinal classification for the families of flowering plants. Annals of the Missouri Botanical Garden. Missouri Botanical Garden 85: 531.

Arseneau, J.-R., R. Steeves, and M. Laflamme. 2017. Modified low-salt CTAB extraction of high-quality DNA from contaminant-rich tissues. Molecular ecology resources 17: 686– 693.

Auguie, B., and A. Antonov. 2017. gridExtra: miscellaneous functions for “grid” graphics. R package version 2.

Avise, J. C., and T. J. Robinson. 2008. Hemiplasy: a new term in the lexicon of phylogenetics. Systematic biology 57: 503–507.

Baker, W. J., P. Bailey, V. Barber, A. Barker, S. Bellot, D. Bishop, L. R. Botigué, et al. 2021. A Comprehensive Phylogenomic Platform for Exploring the Angiosperm Tree of Life. Systematic biology.

Barchi, L., M. Pietrella, L. Venturini, A. Minio, L. Toppino, A. Acquadro, G. Andolfo, et al. 2019. A chromosome-anchored eggplant genome sequence reveals key events in Solanaceae evolution. Scientific reports 9: 11769.

Bennett, J. R. 2008. Revision of *Solanum* section Regmandra (Solanaceae). Edinburgh journal of botany 65: 69–112.

Bohs, L. 2004. A chloroplast DNA phylogeny of *Solanum* section Lasiocarpa. Systematic botany 29: 177–187.

Bohs, L. 2005. Major clades in *Solanum* based on ndhF sequence data. Monographs in Systematic Botany 104: 27.

Bohs, L., and R. G. Olmstead. 2001. A reassessment of *Normania* and *Triguera* (Solanaceae). Plant systematics and evolution 228: 33–48.

Bohs, L., and R. G. Olmstead. 1997. Phylogenetic relationships in *Solanum* (Solanaceae) based on ndhF sequences. Systematic botany 22: 5.

Bolger, A. M., M. Lohse, and B. Usadel. 2014. Trimmomatic: a flexible trimmer for Illumina sequence data. *Bioinformatics (Oxford*, England) 30: 2114–2120.

Bouckaert, R., T. G. Vaughan, J. Barido-Sottani, S. Duchêne, M. Fourment, A. Gavryushkina, J. Heled, et al. 2019. BEAST 2.5: An advanced software platform for Bayesian evolutionary analysis. PLoS computational biology 15: e1006650.

Brown, J. W., J. F. Walker, and S. A. Smith. 2017. Phyx: phylogenetic tools for unix. Bioinformatics 33: 1886–1888.

Capella-Gutiérrez, S., J. M. Silla-Martínez, and T. Gabaldón. 2009. trimAl: a tool for automated alignment trimming in large-scale phylogenetic analyses. *Bioinformatics (Oxford*, England) 25: 1972–1973.

Chavana, J., S. Singh, A. Vazquez, B. Christoffersen, A. Racelis, and R. R. Kariyat. 2021. Local adaptation to continuous mowing makes the noxious weed Solanum elaeagnifolium a superweed candidate by improving fitness and defense traits. Scientific reports 11: 6634.

Chiarini, F., F. Sazatornil, and G. Bernardello. 2018. Data reassessment in a phylogenetic context gives insight into chromosome evolution in the giant genus Solanum (Solanaceae). Systematics and biodiversity 16: 397–416.

Cowie, B. W., N. Venter, E. T. F. Witkowski, M. J. Byrne, and T. Olckers. 2018. A review of *Solanum mauritianum* biocontrol: prospects, promise and problems: a way forward for South Africa and globally. Biocontrol 63: 475–491.

Darriba, D., D. Posada, A. M. Kozlov, A. Stamatakis, B. Morel, and T. Flouri. 2020. ModelTest-NG: A new and scalable tool for the selection of DNA and protein evolutionary models. Molecular biology and evolution 37: 291–294.

Degnan, J. H., and N. A. Rosenberg. 2009. Gene tree discordance, phylogenetic inference and the multispecies coalescent. Trends in ecology & evolution 24: 332–340.

Dodsworth, S., M. J. M. Christenhusz, J. G. Conran, M. S. Guignard, S. Knapp, M. Struebig, A. R. Leitch, and M. W. Chase. 2020. Extensive plastid-nuclear discordance in a recent radiation of *Nicotiana* section Suaveolentes (Solanaceae). Botanical journal of the Linnean Society. Linnean Society of London 193: 546–559.

Duvall, M. R., S. V. Burke, and D. C. Clark. 2020. Plastome phylogenomics of Poaceae: alternate topologies depend on alignment gaps. Botanical journal of the Linnean Society. Linnean Society of London 192: 9–20.

Echeverría-Londoño, S., T. Särkinen, I. S. Fenton, A. Purvis, and S. Knapp. 2020. Dynamism and context-dependency in diversification of the megadiverse plant genus *Solanum* (Solanaceae). Journal of systematics and evolution 58: 767–782.

Edelman, N. B., P. B. Frandsen, M. Miyagi, B. Clavijo, J. Davey, R. B. Dikow, G. García-Accinelli, et al. 2019. Genomic architecture and introgression shape a butterfly radiation. Science 366: 594–599.

Edgar, R. C. 2010. Search and clustering orders of magnitude faster than BLAST. Bioinformatics 26: 2460–2461.

Ewels, P., M. Magnusson, S. Lundin, and M. Käller. 2016. MultiQC: summarize analysis results for multiple tools and samples in a single report. Bioinformatics 32: 3047–3048.

Gardner, E. M., M. Garner, R. Cowan, S. Dodsworth, N. Epitawalage, D. Arifiani, S. Sahromi, et al. 2020. Repeated parallel losses of inflexed stamens in Moraceae: phylogenomics and generic revision of the tribe Moreae and the reinstatement of the tribe Olmedieae (Moraceae). bioRxiv.

Gates, D. J., D. Pilson, and S. D. Smith. 2018. Filtering of target sequence capture individuals facilitates species tree construction in the plant subtribe Iochrominae (Solanaceae). Molecular phylogenetics and evolution 123: 26–34.

Hahn, M. W., and L. Nakhleh. 2016. Irrational exuberance for resolved species trees. Evolution; international journal of organic evolution 70: 7–17.

Hilgenhof, R., E. Gagnon, S. Knapp, X. Aubriot, E.J. Tepe, L. Bohs, L.L. Giacomin, Y.F. Gouvea, J.R. Stehmann, A Orejuela, C.I. Orozco, I.E. Peralta, and T. Sarkinen. 2021. Evolution of key morphological traits in *Solanum* (Solanaceae): identifying areas of interest for further study. American Journal of Botany, in prep.

Hime, P. M., A. R. Lemmon, E. C. M. Lemmon, E. Prendini, J. M. Brown, R. C. Thomson, J. D. Kratovil, et al. 2021. Phylogenomics reveals ancient gene tree discordance in the amphibian tree of life. Systematic biology 70: 49–66.

Huang, B., H. Ruess, Q. Liang, C. Colleoni, and D. M. Spooner. 2019. Analyses of 202 plastid genomes elucidate the phylogeny of *Solanum* section Petota. Scientific reports 9: 4454.

Huson, D. H., and D. Bryant. 2006. Application of phylogenetic networks in evolutionary studies. Molecular biology and evolution 23: 254–267.

Jeffroy, O., H. Brinkmann, F. Delsuc, and H. Philippe. 2006. Phylogenomics: the beginning of incongruence? Trends in genetics: TIG 22: 225–231.

Jin, J.-J., W.-B. Yu, J.-B. Yang, Y. Song, C. W. dePamphilis, T.-S. Yi, and D.-Z. Li. 2020. GetOrganelle: a fast and versatile toolkit for accurate de novo assembly of organelle genomes. Genome biology 21: 241.

Johnson, M. G., E. M. Gardner, Y. Liu, R. Medina, B. Goffinet, A. J. Shaw, N. J. C. Zerega, and N. J. Wickett. 2016. HybPiper: Extracting coding sequence and introns for phylogenetics from high-throughput sequencing reads using target enrichment. Applications in plant sciences 4: 1600016.

Johnson, M. G., L. Pokorny, S. Dodsworth, L. R. Botigué, R. S. Cowan, A. Devault, W. L. Eiserhardt, et al. 2019. A universal probe set for targeted sequencing of 353 nuclear genes from any flowering plant designed using k-Medoids clustering. Systematic biology 68: 594–606.

Junier, T., and E. M. Zdobnov. 2010. The Newick utilities: high-throughput phylogenetic tree processing in the UNIX shell. *Bioinformatics (Oxford*, England) 26: 1669–1670.

Katoh, K., K.-I. Kuma, H. Toh, and T. Miyata. 2005. MAFFT version 5: improvement in accuracy of multiple sequence alignment. Nucleic acids research 33: 511–518.

Knapp, S. 2013. A revision of the Dulcamaroid clade of *Solanum* L. (Solanaceae). PhytoKeys 22: 1–432.

Knapp, S., G. E. Barboza, L. Bohs, and T. Särkinen. 2019. A revision of the Morelloid clade of *Solanum* L. (Solanaceae) in North and Central America and the Caribbean. PhytoKeys 123: 1–144.

Knapp, S., and Others. 2000. A revision of *Solanum thelopodium* species group (section Anthoresis sensu Seithe, pro parte): Solanaceae. *Bulletin of the Natural History Museum*, Botany Series 30: 13–30.

Knapp, S., E. Sagona, A. K. Z. Carbonell, and F. Chiarini. 2017. A revision of the Solanum elaeagnifolium clade (Elaeagnifolium clade; subgenus Leptostemonum, Solanaceae). PhytoKeys: 1–104.

Knapp, S., and M. S. Vorontsova. 2016. A revision of the “African Non-Spiny” Clade of *Solanum* L. (*Solanum* sections Afrosolanum Bitter, Benderianum Bitter, Lemurisolanum Bitter, Lyciosolanum Bitter, Macronesiotes Bitter, and Quadrangulare Bitter: Solanaceae). PhytoKeys 66: 1–142.

Koenen, E. J. M., D. I. Ojeda, and F. T. Bakker. 2021. The origin of the legumes is a complex paleopolyploid phylogenomic tangle closely associated with the cretaceous–paleogene (K–Pg) mass extinction event. Systematic biology 70(3): 508–526.

Kumar, S., A. J. Filipski, F. U. Battistuzzi, S. L. Kosakovsky Pond, and K. Tamura. 2012. Statistics and truth in phylogenomics. Molecular biology and evolution 29: 457–472.

Kuramae, E. E., V. Robert, B. Snel, M. Weiss, and T. Boekhout. 2006. Phylogenomics reveal a robust fungal tree of life. FEMS yeast research 6: 1213–1220.

Lanfear, R., B. Calcott, S. Y. W. Ho, and S. Guindon. 2012. Partitionfinder: combined selection of partitioning schemes and substitution models for phylogenetic analyses. Molecular biology and evolution 29: 1695–1701.

Larridon, I., T. Villaverde, A. R. Zuntini, L. Pokorny, G. E. Brewer, N. Epitawalage, I. Fairlie, et al. 2020. Tackling Rapid Radiations With Targeted Sequencing. Frontiers in plant science 10: 1655.

Levin, R. A., N. R. Myers, and L. Bohs. 2006. Phylogenetic relationships among the “spiny solanums” (*Solanum* subgenus *Leptostemonum*, Solanaceae). American journal of botany 93: 157–169.

Levin, R. A., K. Watson, and L. Bohs. 2005. A four-gene study of evolutionary relationships in *Solanum* section Acanthophora. American journal of botany 92: 603–612.

Li, H., and R. Durbin. 2010. Fast and accurate long-read alignment with Burrows–Wheeler transform. Bioinformatics 26: 589–595.

Liu, L., L. Yu, L. Kubatko, D. K. Pearl, and S. V. Edwards. 2009. Coalescent methods for estimating phylogenetic trees. Molecular phylogenetics and evolution 53: 320–328.

Louca, S., and M. W. Pennell. 2020. Extant timetrees are consistent with a myriad of diversification histories. Nature 580: 502–505.

Lutteropp, S., C. Scornavacca, A. M. Kozlov, and B. Morel. 2021. NetRAX: Accurate and Fast Maximum Likelihood Phylogenetic Network Inference. bioRxiv.

Malinsky, M., H. Svardal, A. M. Tyers, E. A. Miska, M. J. Genner, G. F. Turner, and R. Durbin. 2018. Whole-genome sequences of Malawi cichlids reveal multiple radiations interconnected by gene flow. Nature ecology & evolution 2: 1940–1955.

Mallet, J., N. Besansky, and M. W. Hahn. 2016. How reticulated are species? BioEssays: news and reviews in molecular, cellular and developmental biology.

Meleshko, O., M. D. Martin, T. S. Korneliussen, C. Schröck, P. Lamkowski, J. Schmutz, A. Healey, et al. 2021. Extensive Genome-Wide Phylogenetic Discordance Is Due to Incomplete Lineage Sorting and Not Ongoing Introgression in a Rapidly Radiated Bryophyte Genus. Molecular biology and evolution 38: 2750–2766.

Mendes, F. K., Y. Hahn, and M. W. Hahn. 2016. Gene Tree Discordance Can Generate Patterns of Diminishing Convergence over Time. Molecular biology and evolution 33: 3299– 3307.

Miller, J. S., A. Kamath, and R. A. Levin. 2009. Do multiple tortoises equal a hare? The utility of nine noncoding plastid regions for species-level phylogenetics in tribe Lycieae (Solanaceae). Systematic botany 34: 796–804.

Miller, M. A., W. Pfeiffer, and T. Schwartz. 2010. Creating the CIPRES Science Gateway for inference of large phylogenetic trees. 2010 Gateway Computing Environments Workshop (GCE), IEEE.

Minh, B. Q., M. W. Hahn, and R. Lanfear. 2020. New methods to calculate concordance factors for phylogenomic datasets. Molecular biology and evolution 37: 2727–2733.

Minh, B. Q., H. A. Schmidt, O. Chernomor, D. Schrempf, M. D. Woodhams, A. von Haeseler, and R. Lanfear. 2020. IQ-TREE 2: new models and efficient methods for phylogenetic inference in the genomic era. Molecular biology and evolution 37: 1530–1534.

Morgan, C. C., P. G. Foster, A. E. Webb, D. Pisani, J. O. McInerney, and M. J. O’Connell. 2013. Heterogeneous models place the root of the placental mammal phylogeny. Molecular biology and evolution 30: 2145–2156.

Olave, M., and A. Meyer. 2020. Implementing large genomic single nucleotide polymorphism data sets in phylogenetic network reconstructions: a case study of particularly rapid radiations of cichlid fish. Systematic biology 69: 848–862.

One Thousand Plant Transcriptomes Initiative. 2019. One thousand plant transcriptomes and the phylogenomics of green plants. Nature 574: 679–685.

Paradis, E., and K. Schliep. 2019. ape 5.0: an environment for modern phylogenetics and evolutionary analyses in R. Bioinformatics 35: 526–528.

Parins-Fukuchi, C., G. W. Stull, and S. A. Smith. 2021. Phylogenomic conflict coincides with rapid morphological innovation. Proceedings of the National Academy of Sciences of the United States of America 118.

Pease, J. B., D. C. Haak, M. W. Hahn, and L. C. Moyle. 2016. Phylogenomics reveal three sources of adaptive variation during a rapid radiation. PLoS biology 14: e1002379.

Pellicer, J., and I. J. Leitch. 2020. The Plant DNA C-values database (release 7.1): an updated online repository of plant genome size data for comparative studies. The new phytologist 226: 301–305.

Peralta, I., S. Knapp, and D. Spooner. 2007. The taxonomy of tomatoes: a revision of wild tomatoes (*Solanum* L. section Lycopersicon (Mill.) Wettst.) and their outgroup relatives (*Solanum* sections Juglandifolium (Rydb.) Child and Lycopersicoides (Child) Peralta). Systematic botany monographs 84: 1–186.

Philippe, H., H. Brinkmann, D. V. Lavrov, D. T. J. Littlewood, M. Manuel, G. Wörheide, and D. Baurain. 2011. Resolving difficult phylogenetic questions: why more sequences are not enough. PLoS biology 9: e1000602.

Philippe, H., D. M. de Vienne, V. Ranwez, B. Roure, D. Baurain, and F. Delsuc. 2017. Pitfalls in supermatrix phylogenomics. European Journal of Taxonomy 283. doi: 10.5852/ejt.2017.283

Poczai, P. 2013. To network or not to network, that is the question. Journal of genetics 92: 703– 705.

Price, M. N., P. S. Dehal, and A. P. Arkin. 2010. FastTree 2 – approximately maximum-likelihood trees for large alignments. PloS one 5: e9490.

Rambaut, A., A. J. Drummond, D. Xie, G. Baele, and M. A. Suchard. 2018. Posterior summarization in bayesian phylogenetics using Tracer 1.7. Systematic biology 67: 901– 904.

Reddy, S., R. T. Kimball, A. Pandey, P. A. Hosner, M. J. Braun, S. J. Hackett, K.-L. Han, et al. 2017. Why Do Phylogenomic data sets yield conflicting trees? Data type influences the avian tree of life more than taxon sampling. Systematic biology 66: 857–879.

Romiguier, J., V. Ranwez, F. Delsuc, N. Galtier, and E. J. P. Douzery. 2013. Less is more in mammalian phylogenomics: AT-rich genes minimize tree conflicts and unravel the root of placental mammals. Molecular biology and evolution 30: 2134–2144.

Ronco, F., M. Matschiner, A. Böhne, A. Boila, H. H. Büscher, A. El Taher, A. Indermaur, et al. 2021. Drivers and dynamics of a massive adaptive radiation in cichlid fishes. Nature 589: 76–81.

Rosario, L. H., J. O. Rodríguez Padilla, D. R. Martínez, A. M. Grajales, J. A. Mercado Reyes, G. J. Veintidós Feliu, B. Van Ee, and D. Siritunga. 2019. DNA barcoding of the Solanaceae family in Puerto Rico including endangered and endemic species. Journal of the American Society for Horticultural Science 144: 363–374.

Saarela, J. M., S. V. Burke, W. P. Wysocki, M. D. Barrett, L. G. Clark, J. M. Craine, P. M. Peterson, et al. 2018. A 250 plastome phylogeny of the grass family (Poaceae): topological support under different data partitions. PeerJ 6: e4299.

Sang, T., D. Crawford, and T. Stuessy. 1997. Chloroplast DNA phylogeny, reticulate evolution, and biogeography of Paeonia (Paeoniaceae). American journal of botany 84: 1120.

Särkinen, T., G. E. Barboza, and S. Knapp. 2015. True Black nightshades: phylogeny and delimitation of the Morelloid clade of *Solanum*. Taxon 64: 945–958.

Särkinen, T., L. Bohs, R. G. Olmstead, and S. Knapp. 2013. A phylogenetic framework for evolutionary study of the nightshades (Solanaceae): a dated 1000-tip tree. BMC evolutionary biology 13: 214.

Särkinen, T., P. Poczai, G. E. Barboza, G. M. van der Weerden, M. Baden, and S. Knapp. 2018. A revision of the Old World Black Nightshades (Morelloid clade of *Solanum* L., Solanaceae). PhytoKeys: 1–223.

Sayyari, E., and S. Mirarab. 2016. Fast Coalescent-based computation of local branch support from quartet frequencies. Molecular biology and evolution 33: 1654–1668.

Sayyari, E., and S. Mirarab. 2018. Testing for polytomies in phylogenetic species trees using quartet frequencies. Genes 9.

Schrempf, D., and G. Szöllősi. 2020. The sources of phylogenetic conflicts. Phylogenetics in the Genomic Era: 3.1:1–3.1:23.

Simion, P., H. Philippe, D. Baurain, M. Jager, D. J. Richter, A. Di Franco, B. Roure, et al. 2017. A large and consistent phylogenomic dataset supports sponges as the sister group to all other animals. Current biology 27: 958–967.

Solís-Lemus, C., P. Bastide, and C. Ané. 2017. PhyloNetworks: a package for phylogenetic networks. Molecular biology and evolution 34: 3292–3298.

Spalink, D., K. Stoffel, G. K. Walden, A. M. Hulse-Kemp, T. A. Hill, A. Van Deynze, and L. Bohs. 2018. Comparative transcriptomics and genomic patterns of discordance in Capsiceae (Solanaceae). Molecular phylogenetics and evolution 126: 293–302.

Spooner, D. M., N. Alvarez, I. E. Peralta, and A. M. Clausen. 2016. Taxonomy of wild potatoes and their relatives in Southern South America (*Solanum* sect. Petota and Etuberosum). Systematic botany monographs 100: 1–240.

Spooner, D. M., R. G. van den Berg, A. Rodrigues, J. B. Bamberg, R. J. Hijmans, and S. Lara-Cabrera. 2004. Wild potatoes (*Solanum* section Petota; Solanaceae) of North and Central America. Systematic botany monographs 68: 1–209.

Spooner, D. M., S. Jansky, F. Rodríguez, R. Simon, M. Ames, D. Fajardo, and R. O. Castillo. 2019. Taxonomy of wild potatoes in northern South America (*Solanum* section Petota). Systematic botany monographs 108: 1–305.

Springer, M. S., and J. Gatesy. 2018. Delimiting Coalescence Genes (C-Genes) in Phylogenomic Data Sets. Genes 9.

Stamatakis, A. 2014. RAxML version 8: a tool for phylogenetic analysis and post-analysis of large phylogenies. Bioinformatics 30: 1312–1313.

Stamatakis, A. 2006. RAxML-VI-HPC: maximum likelihood-based phylogenetic analyses with thousands of taxa and mixed models. Bioinformatics 22: 2688–2690.

Stern, S., M. de F. Agra, and L. Bohs. 2011. Molecular delimitation of clades within New World species of the “spiny solanums” (*Solanum* subg. Leptostemonum). Taxon 60: 1429–1441.

Stern, S., and L. Bohs. 2012. An explosive innovation: phylogenetic relationships of *Solanum* section Gonatotrichum (Solanaceae). PhytoKeys: 89–98.

Strickler, S. R., A. Bombarely, J. D. Munkvold, T. York, N. Menda, G. B. Martin, and L. A. Mueller. 2015. Comparative genomics and phylogenetic discordance of cultivated tomato and close wild relatives. PeerJ 3: e793.

Suh, A. 2016. The phylogenomic forest of bird trees contains a hard polytomy at the root of Neoaves. Zoologica scripta 45: 50–62.

Suh, A., L. Smeds, and H. Ellegren. 2015. The dynamics of incomplete lineage sorting across the ancient adaptive radiation of neoavian birds. PLoS biology 13: e1002224.

Symon, D. E. 1994. Kangaroo apples: Solanum sect. Archaesolanum. Adelaide, Australia: Published by the author.

Taberlet, P., L. Gielly, G. Pautou, and J. Bouvet. 1991. Universal primers for amplification of three non-coding regions of chloroplast DNA. Plant molecular biology 17: 1105–1109.

Tavaré, S. 1986. Some probabilistic and statistical problems in the analysis of DNA sequences. Lectures on mathematics in the life sciences 17: 57–86.

Tepe, E. J., G. J. Anderson, D. M. Spooner, and L. Bohs. 2016. Relationships among wild relatives of the tomato, potato, and pepino. Taxon 65: 262–276.

Than, C., D. Ruths, and L. Nakhleh. 2008. PhyloNet: a software package for analyzing and reconstructing reticulate evolutionary relationships. BMC bioinformatics 9: 322.

Tillich, M., P. Lehwark, T. Pellizzer, E. S. Ulbricht-Jones, A. Fischer, R. Bock, and S. Greiner. 2017. GeSeq-versatile and accurate annotation of organelle genomes. Nucleic acids research 45: W6–W11.

Villanueva, R. A. M., and Z. J. Chen. 2019. ggplot2: elegant graphics for data analysis (2nd ed.). Measurement: interdisciplinary research and perspectives 17: 160–167.

Weese, T. L., and L. Bohs. 2007. A three-gene phylogeny of the genus *Solanum* (Solanaceae). Systematic botany 32: 445–463.

Wen, D., Y. Yu, J. Zhu, and L. Nakhleh. 2018. Inferring phylogenetic networks using PhyloNet. Systematic biology 67: 735–740.

Wendel, J. F., and J. J. Doyle. 1998. Phylogenetic incongruence: window into genome history and molecular evolution. *In* D. E. Soltis, P. S. Soltis, and J. J. Doyle [eds.], Molecular Systematics of Plants II: DNA Sequencing, 265–296. Springer US, Boston, MA.

Wickett, N. J., S. Mirarab, N. Nguyen, T. Warnow, E. Carpenter, N. Matasci, S. Ayyampalayam, et al. 2014. Phylotranscriptomic analysis of the origin and early diversification of land plants. Proceedings of the National Academy of Sciences of the United States of America 111: E4859–68.

Wickham, H., and M. H. Wickham. 2019. Package ‘stringr.’

Williams, T. A., D. Schrempf, G. J. Szöllősi, C. J. Cox, P. G. Foster, and T. M. Embley. 2021. Inferring the deep past from molecular data. Genome biology and evolution 13.

Wu, M., J. L. Kostyun, and L. C. Moyle. 2019. Genome sequence of *Jaltomata* addresses rapid reproductive trait evolution and enhances comparative genomics in the hyper-diverse Solanaceae. Genome biology and evolution 11: 335–349.

Yu, G. 2020. Using ggtree to visualize data on tree-like structures. Current protocols in bioinformatics 69: e96.

Zhang, C., M. Rabiee, E. Sayyari, and S. Mirarab. 2018. ASTRAL-III: polynomial time species tree reconstruction from partially resolved gene trees. BMC bioinformatics 19.

